# On prs for complex polygenic trait prediction

**DOI:** 10.1101/447797

**Authors:** Bingxin Zhao, Fei Zou

## Abstract

Polygenic risk score (PRS) is the state-of-art prediction method for complex traits using summary level data from discovery genome-wide association studies (GWAS). The PRS, as its name suggests, is designed for polygenic traits by aggregating small genetic effects from a large number of causal SNPs and thus is viewed as a powerful method for predicting complex polygenic traits by the genetics community. However, one concern is that the prediction accuracy of PRS in practice remains low with little clinical utility, even for highly heritable traits. Another practical concern is whether genome-wide SNPs should be used in constructing PRS or not. To address the two concerns, we investigate PRS both empirically and theoretically. We show how the performance of PRS is influenced by the triplet (*n, p, m*), where *n, p, m* are the sample size, the number of SNPs studied, and the number of true causal SNPs, respectively. For a given heritability, we find that i) when PRS is constructed with all *p* SNPs (referred as GWAS-PRS), its prediction accuracy is controlled by the *p/n* ratio; while ii) when PRS is built with a set of top-ranked SNPs that pass a pre-specified threshold (referred as threshold-PRS), its accuracy varies depending on how sparse the true genetic signals are. Only when *m* is magnitude smaller than *n*, or genetic signals are sparse, can threshold-PRS perform well and outperform GWAS-PRS. Our results demystify the low performance of PRS in predicting highly polygenic traits, which will greatly increase researchers’ awareness of the power and limitations of PRS, and clear up some confusion on the clinical application of PRS.

## 1. Introduction

With the rapid development in biomedical technologies, various types of large-scale genetics and genomics data have been collected for better understanding of genetic etiologies underlying complex human diseases and traits. Genome-wide association studies (GWAS) aim to examine association between complex traits and single-nucleotide polymorphisms (SNPs). To detect SNPs that are associated with a given phenotype, GWAS commonly perform single SNP analysis to estimate and test the association between the phenotype and each candidate SNP one at a time, while adjusting for the effects of non-genetic factors and population sub-structures (Price et al., 2006). Tens of thousands of statistically significant SNPs have been detected for hundreds of human diseases/traits through GWAS (MacArthur et al., 2016; Visscher et al., 2017). However, most of the identified SNPs have very low marginal genetic effects, explaining only a very small portion of the phenotypic variation even for traits with known high heritability (Visscher et al., 2012), resulting in a so called “missing heritability” phenomenon (Manolio et al., 2009; Zuk et al., 2012).

One explanation for the missing heritability is that most complex traits are polygenic, affected by many genes whose individual effects are small (Timpson et al., 2018). The polygenicity of traits has long been hypothesized (Penrose, 1953; Fisher, 1919; Gottesman and Shields, 1967) and is supported by increasing empirical evidence (Yang et al., 2015, 2010; Dudbridge, 2016; Ge et al., 2017; Shi, Kichaev and Pasaniuc, 2016; Lee et al., 2012; Kemp et al., 2017; Wray et al., 2018). Given the polygenicity nature of complex traits, it has been hypothesized that many causal SNPs are not likely to pass the genome-wide significance threshold, but should be informative and used for polygenic trait prediction. For these reasons, the polygenic risk score (Purcell et al., 2009) (PRS) is proposed, as a weighted sum of top-ranked candidate SNPs where each SNP is weighted by its estimated marginal effect from a discovery GWAS. As its name suggests, PRS aims to aggregate genetic effects of polygenes. Thus, it is expected to be powerful for polygenic and omnigenic traits (Boyle, Li and Pritchard, 2017). The omnigenic model is a newly emerging model for complex traits that are affected by the majority (if not all) of candidate SNPs.

In PRS, each SNP is weighted by its GWAS summary statistics, which avoids the need of accessing individual-level genotype data of discovery GWAS, and largely reduces the computational and data storage burdens and bypasses privacy concerns of sharing personal DNA information. As the GWAS summary statistics quickly accumulate in large publicly available data repositories (Watanabe et al., 2018), PRS has been widely used for complex traits in different domains. There were over 3,000 PRS-related publications in 2018. However, the prediction power of PRS remains disappointingly low with little clinical utility, even for traits with known high heritability (Zheutlin and Ross, 2018; Márquez-Luna, Loh and Price, 2017; Torkamani, Wineinger and Topol, 2018). Two legitimate reasons for the poor performance of PRS include 1) low quality SNP arrays with low coverage of causal SNPs; and 2) low quality top-ranked SNPs in tagging causal SNPs (Chatterjee, Shi and García-Closas, 2016; Wray et al., 2013). However, as will be shown by the paper, even in the absence of the above two reasons, PRS may still perform poorly. Thus far, except few studies (Daetwyler, Villanueva and Woolliams, 2008; Dudbridge, 2013; Chatterjee et al., 2013; Vilhjálmsson et al., 2015; Pasaniuc and Price, 2017), little research has been seriously done to study the statistical properties of PRS for polygenic and omnigenic traits.

We aim to fill the gap by empirically and theoretically studying PRS under various polygenic model assumptions with a hope to clear some misperceptions on PRS and to provide some practical guidelines on the use of PRS. Since PRS is built upon the marginal SNP effect estimates, we start our investigation from the statistical properties of the estimates. The performance of PRS is closely related to the asymptotic behavior of sure independent screening (Fan and Lv, 2008) (SIS) when signals are dense. It turns out, our theoretical investigation on the marginal genetic effect estimates is highly relevant to the “spurious correlation” problem (Fan, Guo and Hao, 2012) associated with high and ultra-high dimensional data, which provides another perspective on PRS. Linking the performance of single SNP analysis to the spurious correlation problem makes another significant contribution in understanding the behavior of the single SNP analysis, one of the most commonly used GWAS analysis approaches. For complex polygenic traits, spurious correlation makes the separation of causal and null SNPs difficult (Figure 1). As will be illustrated later, even for a fully heritable trait with a 100% genetic heritability, the estimated genetic effects of causal and null SNPs can be totally mixed and nonseparable from each other. The prediction power of PRS can go as low as zero.

**Figure 1:**
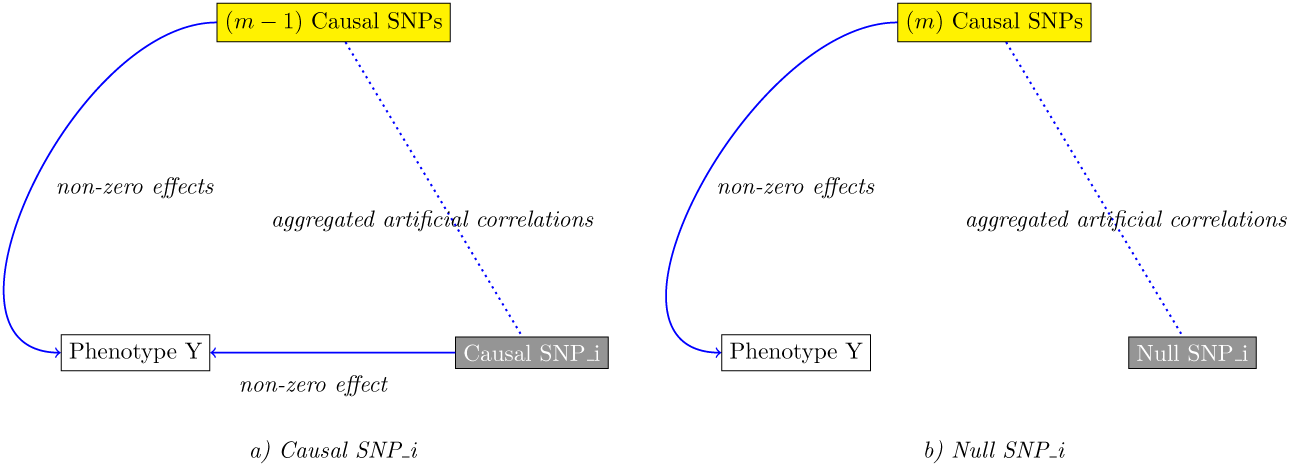
Impact of spurious correlation on the marginal SNP effect estimate of SNP *i* (*i* = 1, …, *p*).

In recognition of the relationship between the GWAS marginal screening and PRS, we show how the asymptotic prediction accuracy of PRS is affected by the triplet of (*n, p, m*), where *n* is the sample size of the training dataset, *p* is the total number of candidate SNPs, and *m* is the number of causal SNPs. Our investigation on PRS starts from the GWAS-PRS and is generalized to the threshold-PRS. Extensive simulations are carried out to evaluate PRS empirically and to evaluate our theoretical results under finite sample settings.

## 2. Single SNP Analysis

For a training dataset, let ***y*** be an *n×*1 phenotypic vector, ***X***_(1)_ denote an *n × m* matrix of the causal SNPs and ***X***_(2)_ donate an *n ×* (*p* − *m*) matrix of the null SNPs, resulting in an *n × p* matrix of all SNPs donated as ***X*** = [***X***_(1)_, ***X***_(2)_] = [***x***_1_,…, ***x***_*m*_, ***x***_*m*+1_, …, ***x***_*p*_]. Columns of ***X*** are assumed to be independent for simplification. Furthermore, column-wise normalization is performed on ***X*** such that each SNP has mean zero and variance one. Define the following condition:

### Condition 1.

*Entries of* ***X*** = [***X***_(1)_, ***X***_(2)_] *are real-valued independent random variables with mean zero, variance one, and a finite eighth order moment*.

The polygenic model assumes the following relationship between ***y*** and ***X***:

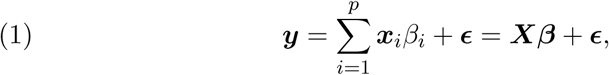

where ***β*** = (*β*_1_, …, *β*_*m*_, *β*_*m*+1_, …, *β*_*p*_)^*T*^ is the vector of SNP effects such that the *β*_*i*_s are i.i.d and follow 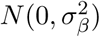 for *i* = 1, …, *m* and *β*_*i*_ = 0 for *i > m*. Let ***β***_(1)_ = (*β*_1_, …, *β*_*m*_)^*T*^ and ***β***_(2)_ be a (*p* − *m*) *×* 1 vector with all elements being zero, and *ϵ* represents the random error vector. For simplicity and without loss of generality, we assume that there exist no other covariates. According to the above model, the overall genetic heritability *h*^2^ of ***y*** is

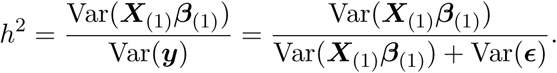

For the rest of the paper, we set *h*^2^ = 1, reducing the above model to the following deterministic model

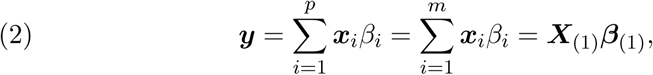

the most optimistic situation in predicting phenotypes.

Though for a typical large-scale discovery GWAS, the sample size *n* is often not small (e.g., *n* ∼ 10, 000 or 100, 000), the number of candidate SNPs *p* is typically even larger (e.g., *p* ∼ 500, 000 or 10, 000, 000). On the other hand, depending on their underlying genetic architectures, the number of causal SNPs can vary dramatically from one trait to another. To cover most of modern GWAS data, we therefore assume *n, p* → ∞ and that

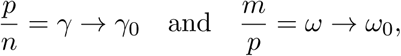

where 0 *< γ*_0_ ≤ ∞, and 0 ≤ *ω*_0_ ≤ 1.

For continuous traits in absence of any covariates, a typical single SNP analysis employs the following simple linear regression model

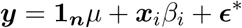

for a given SNP *i*, where **1**_***n***_ is an *n ×* 1 vector of ones, *β*_*i*_ is its effect (*i* = 1, …, *p*). When both ***y*** and ***x***_*i*_ are normalized and *n* → ∞, under Condition (1) and polygenic model (2), the maximum likelihood estimate (MLE) of 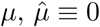, and the MLE of the genetic effect *β*_*i*_ equals

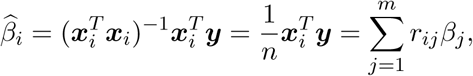

where 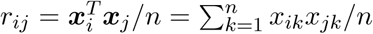 is the sample correlation between ***x***_*i*_ and ***x***_*j*_, *j* = 1,…, *p*. Specifically, for SNP *i, i* = 1, …, *p*, we have

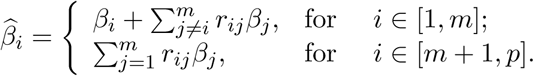

Given that SNPs in ***X*** are independent of each other, or the correlation *ρ*_*ij*_ for any *ij*th SNP pair (*i* & *j*)(*i* ≠ *j*) is zero, it can be shown that asymptotically 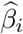 is an unbiased estimator of *β*_*i*_ such that

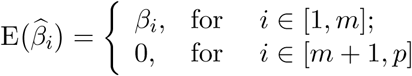

as *n* → ∞. The associated variance of 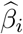 grows linearly with *m* since for any causal SNP *i* (1 ≤ *i* ≤ *m*), we have

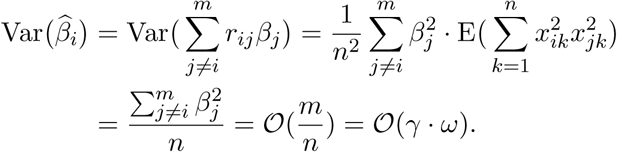

Similarly, for any null SNP *i* (*m < i* ≤ *p*), we have

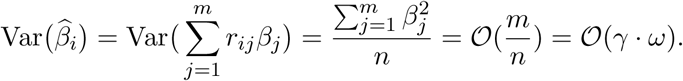

Therefore, 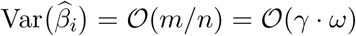 for all *i* (*i* = 1,…, *p*), suggesting that the associated variances of all 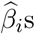 are in the same scale regardless of whether their corresponding SNPs are causal or not. When *m/n* = *γ* · *ω* → *γ*_0_ · *ω*_0_ is large, 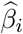 is no longer a reliable estimate of *β*_*i*_. Moreover, the 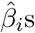 of the causal and null SNPs become well mixed and cannot be easily separated when *m/n* is large, which raises two important concerns include 1) how does this affect the SNP selection; and 2) how does this affect the weights in PRS which ultimately affect the performance of PRS? We address these two concerns below.

## 3. PRS

For a testing dataset with *n*_*z*_ samples, define its *n*_*z*_*×p* SNP matrix as ***Z*** = [***Z***_(1)_, ***Z***_(2)_] with ***Z***_(1)_ = [***z***_1_,…, ***z***_*m*_] and ***Z***_(2)_ = [***z***_*m*+1_,…, ***z***_*p*_]. Then 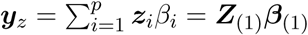 and the PRS equals

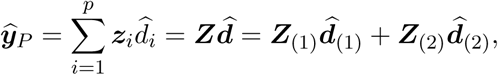

where 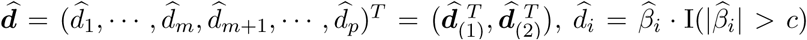, I(·) is the indicator function, and *c* is a given threshold for screening SNPs. When *c* = 0, all candidate SNPs are used, leading to GWAS-PRS. The prediction accuracy of PRS is typically measured by

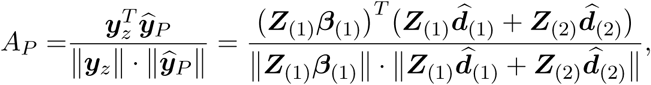

where ‖ · ‖ is the *l*_2_ norm of a vector.

### 3.1 GWAS-PRS

In GWAS-PRS, 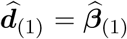 and 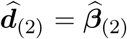. For simplification, in rest of the paper, we set *n*_*z*_ = *n* and our general conclusions should remain the same when the two are different. Let 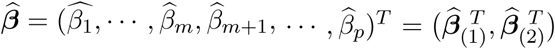, where 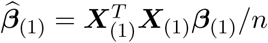, and 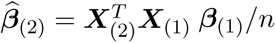 Then

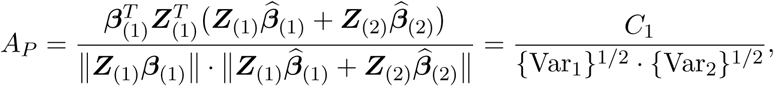

where

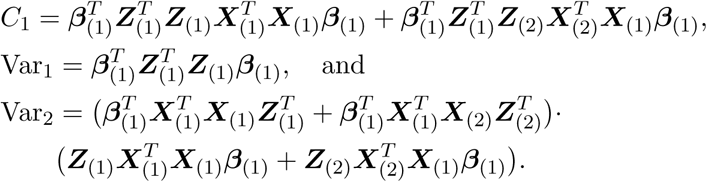

#### Theorem 1.

*Under the polygenic model (2) and Condition (1), if m* → ∞, *p* → ∞, *and p/n*^2^ → 0 *as n* → ∞,

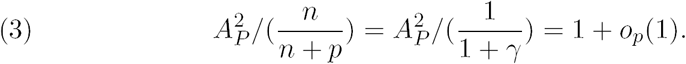

*If we further assume p* = *c·n*^*α*^ *for some constants c* ∈ (0, ∞) *and α* ∈ (0, ∞], *then*

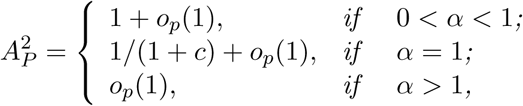

*as n* → ∞.

Proof of Theorem 1 is given in Appendix A.

#### Remark 1.

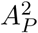 *has nonzero asymptotic limit provided that α* ∈ (0, 1]. *As illustrated in Figure 2, A*_*P*_ *converges to zero if α* ∈ [2, ∞], *indicating the null prediction power of GWAS-PRS even for traits that are fully heritable*.

**Figure 2:**
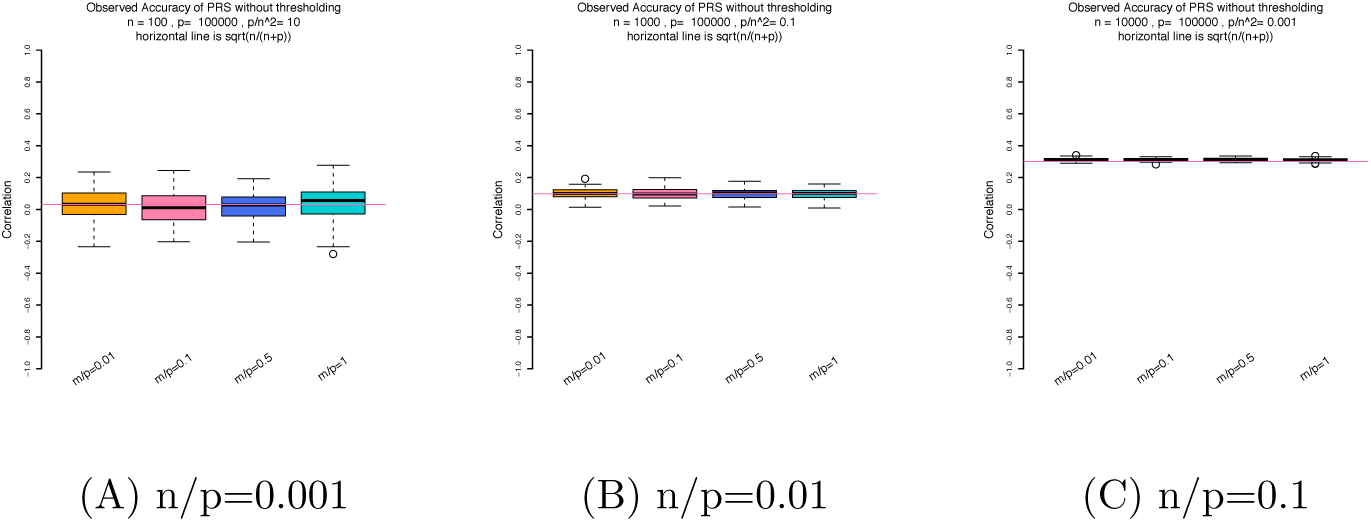
Prediction accuracy (*A*_*P*_) of GWAS-PRS across different *m/p* ratios when *c* = 0 (i.e., all SNPs are selected). We set *p*=100, 000 and *n*=100, 1000 and 10, 000, respectively.

#### Remark 2.

*Since causal SNPs are not known as a prior but estimated, a poorly selected SNP list can be a reason for the poor performance of PRS. However, Theorem 1 suggests that when m is large, even in the oracle situation where the selected SNP list contains all and only all of the causal SNPs, its performance is limited by the ratio of m/n. For complex traits underlying the omnigenic model (Boyle, Li and Pritchard, 2017), the expected prediction power of the oracle PRS is essentially zero*.

#### Remark 3.

*In our investigation, we set h*^2^ = 1 *to reflect the most optimistic situation for phenotype prediction. The result in (3) can be easily extended to* 0 *< h*^2^ *<* 1 *as follows:*

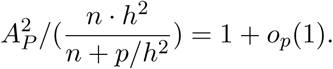

### 3.2 Illustration of asymptotic limits of GWAS-PRS

We numerically evaluate the analytical results above and the performance of 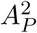 with *p* = 100, 000, and *n* = *n*_*z*_ = 100, 1000 or 10, 000. Each entry of ***X*** and ***Z*** is independently generated from *N* (0, 1). We also vary the ratio of causal SNPs *m/p* from 0.01 to 1 to reflect a wide range of SNP signals, from very sparse to very dense situations, respectively. The effects of the causal SNPs ***β***_(1)_ ∼ *MV N* (0, ***I***_*m*_), based on which the phenotypes ***y*** and ***y***_*z*_ are generated from model (2), where ***I***_*m*_ is an *m × m* identity matrix. A total of 100 replications are conducted for each simulation setup.

Figure 2 shows the *A*_*P*_ values across different simulation setups. As expected, for GWAS-PRS, *A*_*P*_ remains nearly constant regardless of *m* and is close to 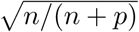. For small *n*, the *A*_*P*_ values are fairly close to zero with a large variance.

### 3.3 Threshold-PRS

As shown in Theorem 1, the asymptotic limit of *A*^2^ associated with GWAS-PRS does not depend on *m*, the number of causal SNPs, but *n*, the sample size of the training dataset. Where sample size is much smaller than the number of candidate SNPs, GWAS-PRS has low prediction accuracy. It is thus natural for us to turn around and ask with a properly selected threshold *c*, whether the performance of PRS can be improved or not.

#### 3.3.1 General Setup

For a given threshold *c >* 0, let *q* = *p*·*a* (*a* ∈ (0, 1]), where *q* is the number of selected SNPs, among which there are *q*_1_ true causal SNPs and the remaining *q*_2_ are null SNPs, that is, *q* = *q*_1_ + *q*_2_. Let ***Z***_(1)_ = [***Z***_(11)_, ***Z***_(12)_], ***Z***_(2)_ = [***Z***_(21)_, ***Z***_(22)_], ***X***_(1)_ = [***X***_(11)_, ***X***_(12)_], 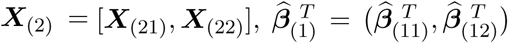 and 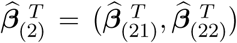, where ***Z***_(11)_, ***X***_(11)_, and 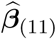 correspond to the selected *q*_1_ causal SNPs, and ***Z***_(21)_, ***X***_(21)_, and 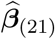 correspond to the selected *q*_2_ null SNPs. The prediction accuracy of threshold-PRS is therefore

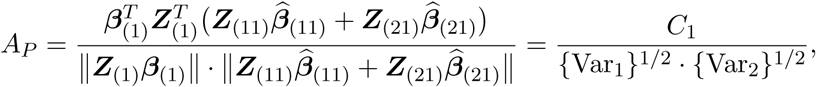

where

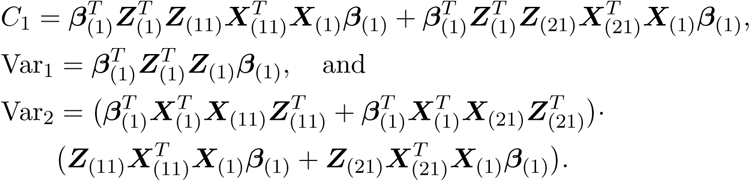

##### Theorem 2.

*Under the polygenic model (2) and Condition (1), if m, q*_1_, *q*_2_ → ∞ *as min*(*n, p*) → ∞, *and further if* 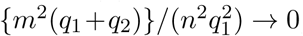, *then we have*

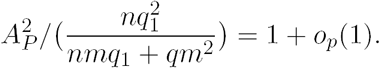

*However, if* 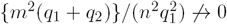 *as min*(*n, p*) → ∞, *then*

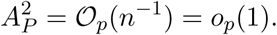

Proof of Theorem 2 is given in Appendix A. Theorem 2 shows that given *n* and *m, A*_*P*_ is determined by *q*_1_, the number of selected causal SNPs, and *q*, the total number of selected SNPs. Expressing *q*_1_ as a function of *q*, or *q*_1_ = *ϕ*(*q*), then *A*_*P*_ can be re-expressed as a function of *q*:

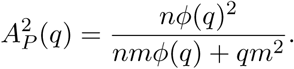

#### 3.3.2 Role of ϕ(*q*)

The function *ϕ*(*q*) is a non-decreasing function of *q* and plays an important role in determining *A*_*P*_. The exact form of *ϕ*(*q*) is trait-dependent and not easy to obtain. But for the following two special examples, we can evaluate *ϕ*(*q*) straightforwardly and investigate its impact on *A*_*P*_. We first study the marginal distribution of the 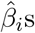, which is a mixture of two distributions, one corresponding to the causal SNP set and one to the null SNP set. Let 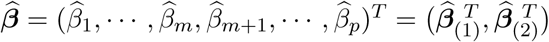. Given that *β*_*i*_s in ***β***_(1)_ are i.i.d and follow 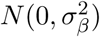, and the remaining ones in ***β***_(2)_ are all zeros, it follows from the central limit theorem that

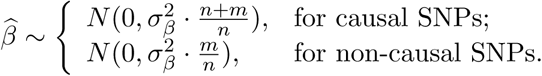

When *m/n* = *γ* · *ω* → *γ*_0_ *· ω*_0_ = 0, the spread of the marginal distribution of the causal SNPs is much wider than that of the marginal distribution of the null SNPs, making the two distributions separable and single SNP analysis powerful. However, as the genetic signal gets denser and denser (or as *m* increases), the difference between the two distributions gets smaller and smaller, leading to two well mixed distributions and poorly performed single SNP analysis. To see how the ratio of *m/n* impacts single SNP analysis, we first approximate the density of 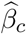 by

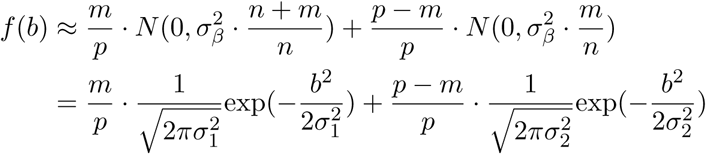

with cumulative distribution function (CDF) given by

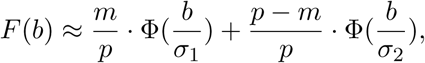

where 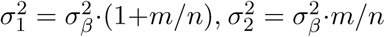, and 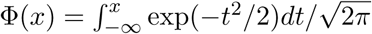 is the CDF of the standard normal random variable. Since the mixture distribution is symmetric about zero, without loss of generality, we consider one-sided test and SNPs with the largest (100*×a*)% estimated genetic effects (0 *< a <* 1*/*2) are selected. For a causal SNP, its selection probability *κ*_1_ equals to

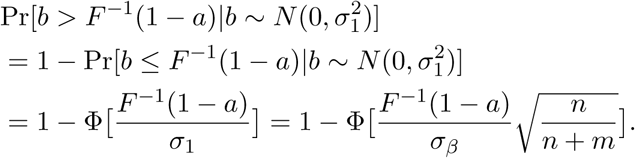

Similarly for a given null SNP, its selection probability *κ*_2_ is

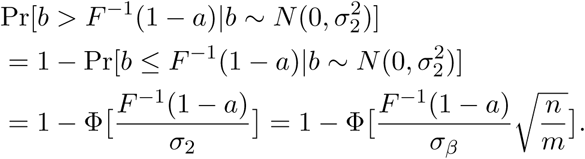

Therefore, among *q* = *p* · *a* = *q*_1_ + *q*_2_ selected SNPs, we have

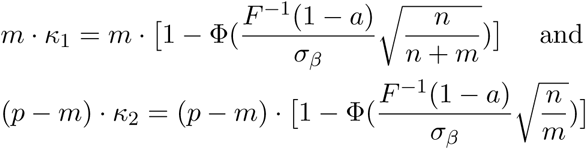

causal and null SNPs, respectively. For a given *a* or equivalently *c, F* ^−1^(1 − *a*)*/σ*_*β*_ is the same for both causal and null SNPs. Therefore, the quality of top-ranked SNP list is largely determined by *m/n*. Remark 4 discusses the upper bounds of *A*_*p*_ under two extreme cases.

##### Remark 4.

*When n/m* = *o*(1), *it is easy to see κ*_1_ = *κ*_2_ · {1 + *o*(1)}. *Therefore, when both q*_1_ *and q*_2_ *are large, q*_1_*/q* ≈ *m/p, and thus* 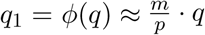, *yielding*

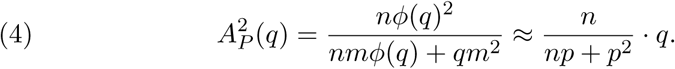

*Therefore, A*_*P*_ *reaches its upper bound* 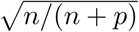, *suggesting that the best performing PRS is the one without SNP selection or GWAS-PRS when the genetic signals are dense. On the other hand, when m/n* = *o*(1), *κ*_1_ *becomes much larger than κ*_2_. *Thus, causal SNPs can be relatively easy to detect by using single SNP analysis. As a increases, q*_1_ *eventually gets saturated at m, and threshold-PRS reaches its upper performance limit* 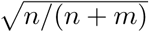 *with q* = *q*_1_ = *ϕ*(*q*) = *m, the oracle case described in Remark 2*.

In conclusion, the above analysis provides some practical guidelines on the use of PRS: 1) SNP screening should be avoided for highly polygenic or omnigenic traits where the *m/n* ratio is large; and 2) for monogenic and oligogenic traits (Timpson et al., 2018) with a small *m/n* ratio, threshold-PRS should be used.

## 4. More simulation studies

### 4.1 Threshold-PRS

To illustrate the finite sample pattern of threshold-PRS, we simulate *p* = 100, 000 uncorrelated SNPs. As in Figure 2, we (naively) generate SNPs from *N* (0, 1). To study the effect of *m/p*, we vary the number of causal SNPs *m* and set it to 100, 1000, 10, 000 and 50, 000. The casual SNP effects ***β***_(1)_ ∼ *MV N* (0, ***I***_*m*_). The linear polygenic model (2) is used to generate phenotype ***y*.** The sample size is set to 1000 and 10, 000 for training data, and 1000 for testing data. For threshold-PRS, we consider a series of *P* -value thresholds {1, 0.8, 0.5, 0.4, 0.3, 0.2, 0.1, 0.08, 0.05, 0.02, 0.01, 10^−3^, 10^−4^, 10^−5^, 10^−6^, 10^−7^, 10^−8^} (Márquez-Luna, Loh and Price, 2017). We name this simulation setting Case 1. Again, a total of 100 replications are conducted for each simulation condition.

Figure 3 and Supplementary Figure 1 display the performance of threshold-PRS across a series of *m/p* ratios in Case 1. As expected, the performance of GWAS-PRS (i.e. at *P* -value threshold of 1) stays nearly constant around 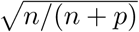 regardless of the *m/p* ratios, which is about 0.3 (shown in Figure 3) for *n* of 10, 000 and 0.1 (shown in Supplementary Figure 1) for *n* of 1000. In contrast, the performance of threshold-PRS varies with *m*. When *m* is small compared to *n*, threshold-PRS performs significantly better than GWAS-PRS provided a reasonable *c* is chosen, which in general is small as shown by Supplementary Figure 1. For example, when *m* = 100 and *n* = 1000, threshold-PRS achieves its best performance at *c* = 10^−5^, with *A*_*P*_ of 0.75, in contrast to its oracle performance which is about 0.95. Figure 3 shows that when *m* gets close to *n* or larger than *n*, the performance of threshold-PRS drops significantly regardless of *c*. When *m* gets close to *n*, its performance remains similar for a wide range of *c* values; and when *m* gets much larger than *n*, its performance improves as *c* increases, and eventually reaches the performance level of GWAS-PRS.

**Figure 3:**
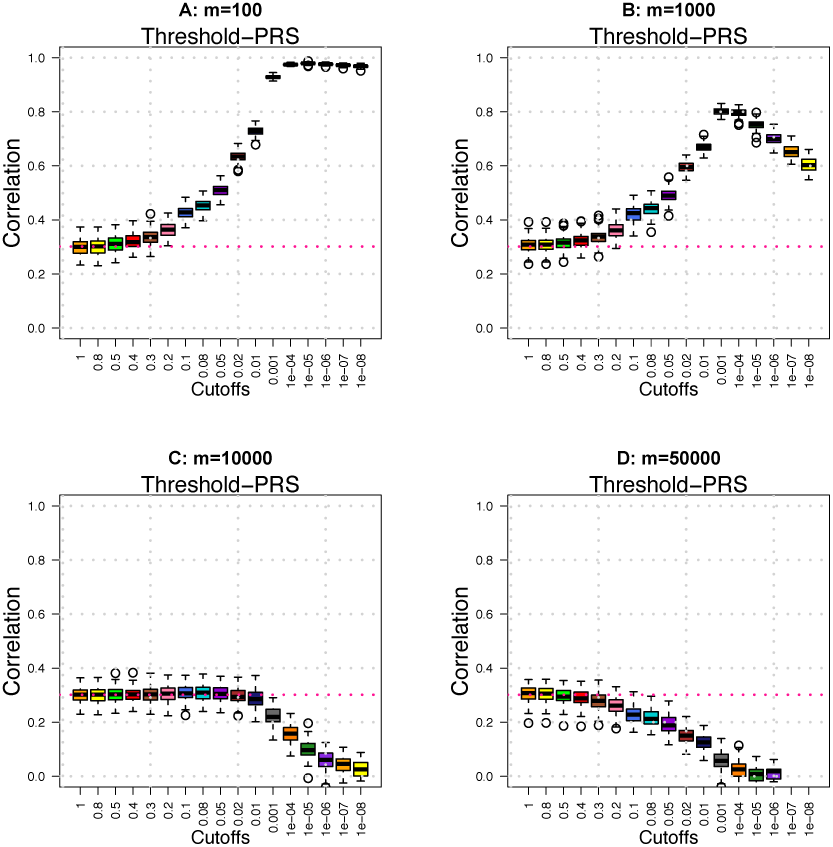
Prediction accuracy (*A*_*P*_) of threshold-PRS across different *m/p* ratios. We set *p*=100, 000 and *n* = 10, 000 in training data and *n*=1000 in testing data.

In addition, we vary Case 1 settings to check the sensitivity of our results. In Case 2, we generate actual SNP genotype data where the minor allele frequency (MAF) of each SNP, *f*, is independently generated from Uniform [0.05, 0.45] and SNP genotypes are independently sampled from {0, 1, 2} with probabilities {(1 − *f*)^2^, 2*f* (1 − *f*), *f* ^2^}, respectively according to the Hardy-Weinberg equilibrium principle. In Case 3, we simulate mixed samples from five subpopulations. The overall MAF of each SNP in mixed samples is then correspondingly generated from Uniform [0.05, 0.45], and the *F*_*st*_ values are independently generated from Uniform [0.01, 0.04] (Lee, Wright and Zou, 2011), based on which the MAF of each sub-population is generated according to the Balding-Nichols model (Balding and Nichols, 1995). We let the sample size of sub-population be the same, and set them at either 200 or 2000. The population substructures are estimated with the principal component analysis (Price et al., 2006) and the top 4 principal components are included as covariates in the single SNP analysis. Case 4 allows larger variability in the causal SNP effects such that *β*_*i*_s are independently generated from 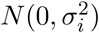, where 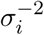 follows a gamma distribution with *α* = 10 and *β* = 9.

The results of Case 2 are displayed in Supplementary Figures 2 and 3, which are similar to those of Case 1. Supplementary Figures 4 and 5 display the oracle performance of PRS under varying *m/p* ratios in Case 2. Clearly *A*_*p*_ is around 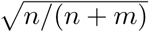, confirming the poor performance of PRS even in the oracle case when genetic signals are dense. The performance of threshold-PRS under Case 2 for traits with *h*^2^=0.5 is presented in Supplementary Figures 6 and 7. Compared to Supplementary Figures 2 and 3 where *h*^2^=1, the prediction accuracy of threshold-PRS decreases, but the general patterns remain the same. The results of Case 3 are displayed in Supplementary Figures 8 and 9. In the presence of population substructures, if they are properly adjusted, the main pattern of threshold-PRS remains unchanged and the performance of GWAS-PRS agrees well with the theoretical results. The results of Case 4 are displayed in Supplementary Figures 10 and 11, which are also similar to those of Case 1, indicating that our asymptotic results are not sensitive to the distribution of the causal SNP effects, or ***β***_(1)_.

### 4.2. UK Biobank SNP data

We perform additional simulation based on real SNP data from the UK Biobank (UKB) resources (Sudlow et al., 2015; Bycroft et al., 2018). We download the unimputed genotype data released in July 2017 and apply the following quality control (QC) procedures: excluding subjects with more than 10% missing genotypes, only including SNPs with MAF > 0.01, genotyping rate > 90%, and passing Hardy-Weinberg test (*P* -value > 1 × 10^−7^). There are 461, 488 SNPs left for 488, 371 subjects after QC, and we randomly select 11, 000 individuals of British ancestry to perform this simulation, among which 10, 000 are randomly selected and used as training data, and PRS are constructed on the remaining 1, 000 individuals. Causal SNPs are randomly selected, and the number of causal SNPs *m* is set to 470, 4700, 47, 000, and 235, 000. The nonzero SNP effects are independently generated from *N* (0, 1), and the heritability *h*^2^ is set to 80%. Single SNP analysis is performed using the PLINK toolset (Purcell et al., 2007). To obtain a list of independent SNPs for PRS construction, we perform linkage disequilibrium (LD)-based SNP clumping for the GWAS results via PLINK. With the default window size (250 kb), we vary the clumping parameter *R*^2^ and set it to 0.05, 0.1, 0.2, 0.3, 0.5, and 0.9. Smaller *R*^2^ results in more stringent selection and more filtered SNPs by clumping. The clumped SNPs are used to construct PRS on testing data.

The results are displayed in Supplementary Figures 12-17, which show similar patterns as in Figure 3. Specifically, GWAS-PRS has consistent performance across the *m/p* ratio and the clumping parameter *R*^2^. The prediction accuracy is around 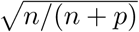, where *p* is the number of clumped SNPs given *R*^2^ = 0.05 (*p* ≈ 150, 000). In addition, threshold-PRS may have high prediction accuracy only when the genetic signals are sparse.

## 5. PRS on Brain Size

We present a real data example to predict the human total brain volume (TBV) with PRS. TBV is generated from brain magnetic resonance imaging (MRI) via advanced normalization tools (Avants et al., 2011) (ANTs) to measure brain size. The UKB sample has 19, 629 participants with both TBV and imputed SNP data (*p* = 8, 944, 375) available after quality controls (Zhao et al., 2019).

We perform a ten-fold analysis where nine folds of the data are used to construct the summary statistics and the remaining one fold of the data is used to measure the PRS prediction accuracy. Twelve PRS are created with p-value thresholds 1, 0.8, 0.5, 0.4, 0.3, 0.2, 0.1, 0.08, 0.05, 0.02, 0.01, and 10^−3^. Next, we used the GWAS summary statistics estimated from all 19, 629 UKB individuals (Zhao et al. (2019), available at https://med.sites.unc.edu/bigs2/data/gwas-summary-statistics/) to construct PRS on subjects from three independent studies, including the Human Connectome Project (HCP, *n* = 1141) study, the Pediatric Imaging, Neurocognition, and Genetics (PING, *n* = 924) study, and the Alzheimer’s Disease Neuroimaging Initiative (ADNI, *n* = 1248) study. See supplementary material for more information, such as SNP data quality controls, SNP pruning and demographic information of these studies.

The prediction accuracy is measured by the (partial) *R*^2^ of PRS from the linear regression model for TBV where the effects of age and gender are adjusted for. Figure 4 displays the average out-of-UKB *R*^2^ across HCP, ADNI, and PING, and the *R*^2^ of within-UKB ten-fold prediction. GWAS-PRS constructed with all candidate SNPs have consistent *R*^2^ in within-UKB ten-fold prediction and out-of-UKB prediction, which is aound 1.5% (*P* - value < 1.92 × 10^−6^). For threshold-PRS, there is a similar pattern that *R*^2^ decreases as the *P* -value threshold decreases, especially when the cutoff is less than 0.2. Such pattern matches with our asymptotic results of threshold-PRS when SNP signals are dense, clearly suggesting that TBV is a complex polygenic trait. We therefore recommend GWAS-PRS for TBV prediction.

**Figure 4:**
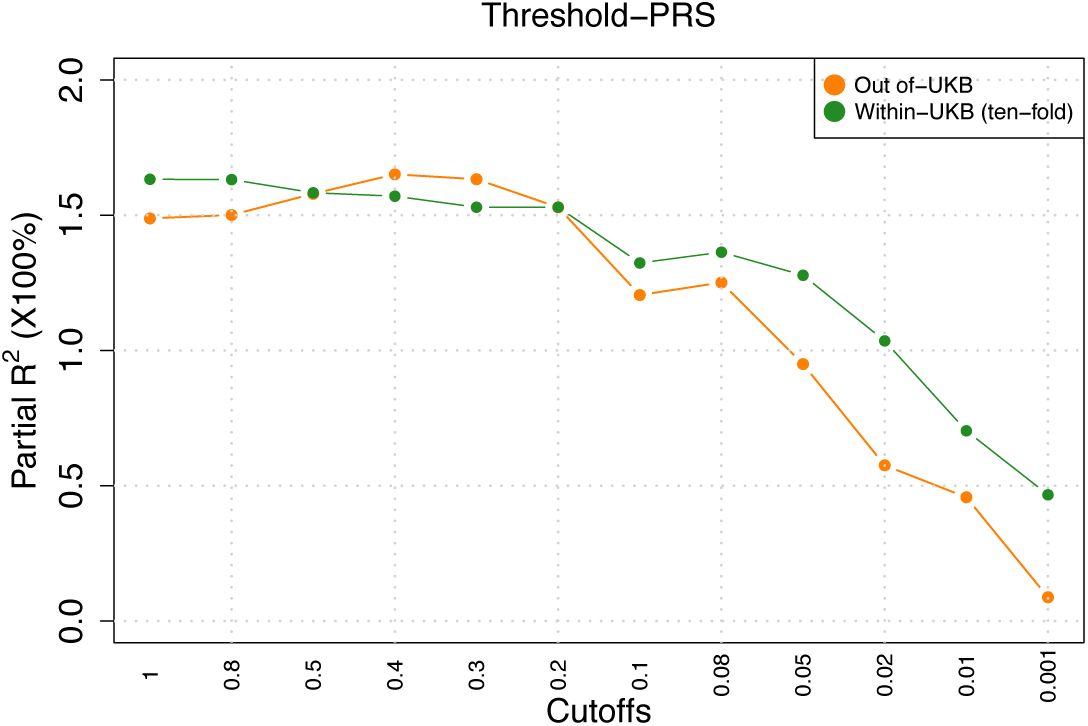
Prediction accuracy (*A*_*P*_) of PRS for total brain volume in ten-fold UKB data prediction and out-of-UKB prediction.

## 6. Discussion

The PRS is widely used in GWAS for predicting phenotypes and for studying co-heritability of pleiotropy phenotypes (Purcell et al., 2009; Chatterjee, Shi and García-Closas, 2016; Power et al., 2015; Choi, Mak and O’Reilly, 2018; Khera et al., 2018). The purpose of PRS is to aggregate effects from a large number of causal SNPs, each of which has small contribution to the phenotype. However, recent years have witnessed widespread not so satisfying performances of PRS in real data applications (Bogdan, Baranger and Agrawal, 2018; Zheutlin and Ross, 2018; Márquez-Luna, Loh and Price, 2017; Torkamani, Wineinger and Topol, 2018; Clarke et al., 2016; Mistry et al., 2018a, b; Socrates et al., 2017). Though previous studies (Daetwyler, Villanueva and Woolliams, 2008; Dudbridge, 2013; Chatterjee et al., 2013) have found that the prediction accuracy of PRS is related to (*n, p*), the asymptotic properties of PRS is largely unknown. More rigorous investigation done in this paper would greatly help researchers gain better insights on PRS and appreciate the limitations of PRS to avoid over- or under-interpreting their findings. In addition, there exists no theoretical study on how the sparsity of genetic signals affects PRS.

We illustrate for the first time how and why the commonly used marginal screening approaches for polygenic traits may fail in preserving the rank of GWAS signals. The properties of single SNP analysis is closely related to the increasingly recognized spurious correlation problem (Fan, Guo and Hao, 2012; Chen, Fan and Li, 2018; Cai et al., 2011; Cai, Fan and Jiang, 2013; Fan et al., 2018; Fan and Zhou, 2016; Su, 2018) in the statistics community. For polygenic/omnigenic traits, single SNP analysis is always misspecified since the effects of a large number of causal SNPs are not modeled but absorbed into the error term, which can profoundly affect the accuracy of marginal screening, an issue that can be safely ignored for traits with sparse genetic signals.

For GWAS-PRS, our asymptotic results align well with previous studies in genetics community (Daetwyler, Villanueva and Woolliams, 2008; Dudbridge, 2013; Chatterjee et al., 2013), but are more statistically rigorous and general. In addition, we illustrate in Figure 2 that the variation of the PRS performance of *A*_*P*_ increases with the increase of the *p/n* ratio as well. When the sample size of training data is small, or *p/n* is large, the prediction accuracy of GWAS-PRS fluctuates around zero with a large variation and the results of PRS should be interpreted with caution. We further generalize our results to threshold-PRS. We highlight how the number of causal SNPs influences the performance of threshold-PRS, and recognize its distinct behaviors under dense and sparse genetic signal scenarios. Threshold-PRS can be powerful for complex traits with sparse genetic signals provided that proper threshold is used. For traits with highly polygenic or omnigenic genetic structures, the general rule is to avoid the use of threshold-PRS but GWAS-PRS. However, the performance of the later is solely determined by the *n/p* ratio and thus is low when *n* is not large, leading to a practical paradox. Further, in our theoretical investigation and simulation studies, we assume all causal SNPs are observed, an assumption that likely not holds for any real GWAS. For real GWAS data with partially genotyped causal SNPs, GWAS-PRS and threshold-PRS are expected to perform even worse.

Last, it is worth mentioning that the limited power of PRS for predicting highly polygenic traits is not unique to PRS, but due to the fundamental analytic challenge in studying such traits. Performed on individual-level data, popular regularization-based methods (such as LASSO (Tibshirani, 1996) or Ridge regression (Hoerl and Kennard, 1970; Tikhonov, 1963)) and the best linear unbiased prediction (BLUP) methods (Chen et al., 2015; Yang et al., 2011) have low prediction power similar to that of PRS unless the genetic signals are spare where LASSO is clearly advantageous. See simulation results in Supplementary Figures 18-20 with simulation details in the supplementary material.

In conclusion, this study thoroughly investigates the power and limitations of PRS on predicting highly polygenic traits. The research hopefully will increase researchers’ awareness on the challenges in studying complex polygenic traits, and the statistics and genetics communities’ awareness on the need of developing novel experiments and innovative statistical methods for complex polygenic traits that are inherently difficult to study.

## Acknowledgements

We would like to thank Ziliang Zhu, Jingwen Zhang, and Hongtu Zhu for helpful discussions; and Xinlei Mi for independent technical verification. We are deeply grateful to Hongtu Zhu for providing helpful comments and edits; and to Laura Zhou for careful proofreading and feedback. The manuscript has been greatly improved by their generous contributions. This research has been conducted using the UK Biobank resource (application number 22783), subject to a data transfer agreement. We thank Hongtu Zhu, Tengfei Li and other members of the UNC BIG-S2 lab for providing and processing the raw imaging data. We thank the individuals represented in the UK Biobank, PING, HCP, and ADNI studies for their participation and the research teams for their work in collecting, processing and disseminating these datasets for analysis. More information of these studies can be found in the supplementary material.

## APPENDIX A: PROOF

In this appendix, we highlight the key steps and important intermediate results to prove the two theorems. More technical details can be found in the supplementary material.

### Proposition A1.

*Under the polygenic model (2) and Condition (1), if m* → ∞ *when n, p* → ∞, *then we have*

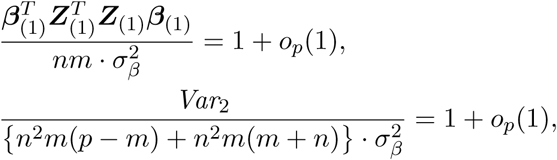

*where*

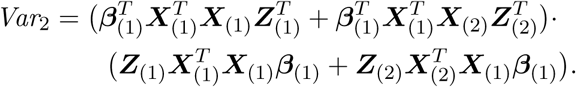

*Further if p/n*^2^ → 0, *then we have*

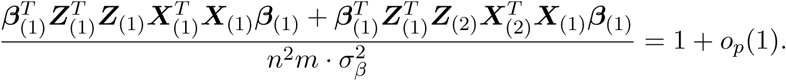

By continuous mapping theorem, we have

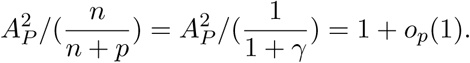

It follows that Theorem 1 is proved for *α* ∈ (0, 2). Now consider the case that *p/n*^2^ ↛ *o*(1), i.e., *α* ∈ [2, ∞]. Note that

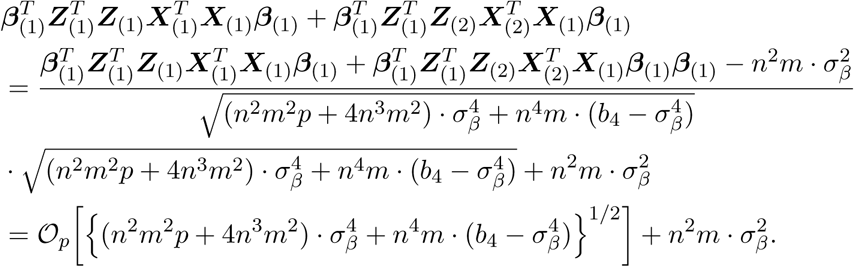

It follows that

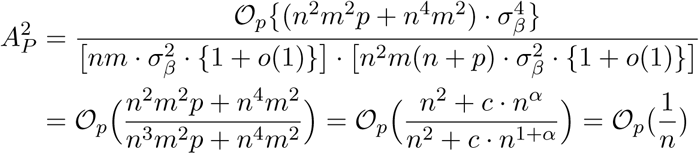

for *α* ∈ [2, ∞]. Thus, Theorem 1 is proved for *α* ∈ (0, ∞].

### Proposition A2.

*Under the polygenic model (2) and Condition (1), if m, q*_1_, *q*_2_ → ∞ *when n, p* → ∞, *then we have*

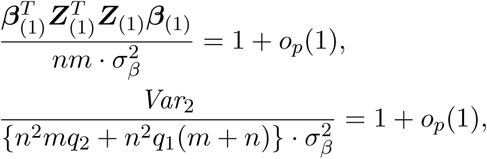

*where*

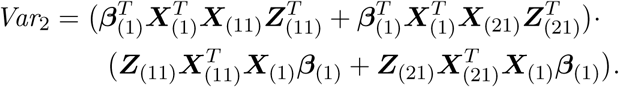

*Further if* 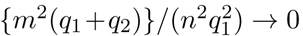, *then we have*

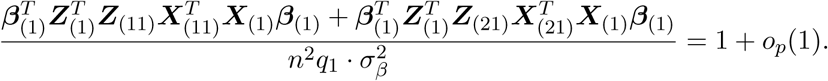

By continuous mapping theorem, we have

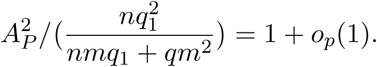

Without loss of generality and for simplicity, we let 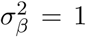, and we note that

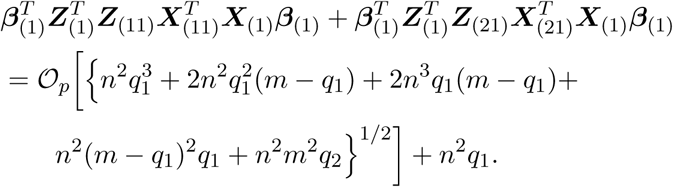

Then if 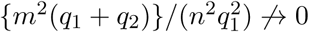, we have

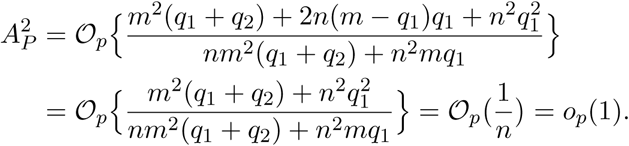

Thus, Theorem 2 is proved.

## SUPPLEMENTARY MATERIAL

**Supplement to: “On PRS for Complex Polygenic Trait Prediction”** (doi: 10.1214/00-AOASXXXXSUPP). We provide additional theoretical de-rails, real data information and simulation results.

**SUPPLEMENT TO “ON PRS FOR COMPLEX POLYGENIC TRAIT PREDICTION”**

## 1. Intermediate results

Proposition S1. *Under Condition (1), if m* → ∞ *as n, p* → ∞, *then we have*

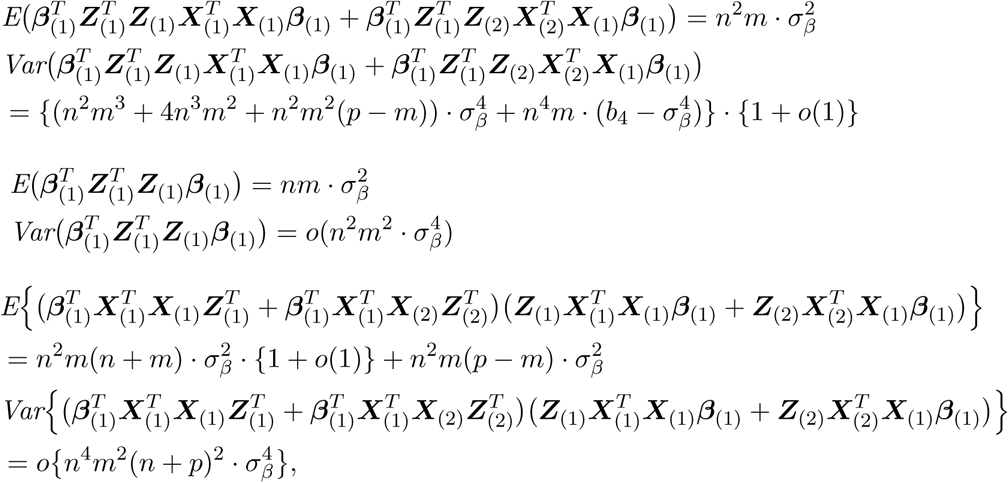

*where b*_4_ < ∞ *is the forth moment of β*.

Proposition S1 quantifies the scale of the three terms in *A*_*P*_. Particularly, for the two variance terms 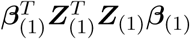 and

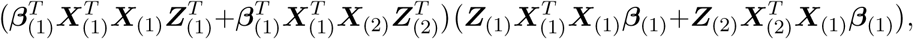

the expected values can respectively dominate the corresponding standard error for any ratios among (*p, m, n*). However, for the covariance term

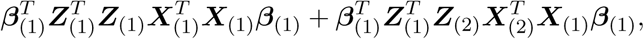

its standard error may or may be dominated by its expected value depending on *p/n*. Following Proposition S1, by Markov’s inequality, for any constant *k* > 0, we have

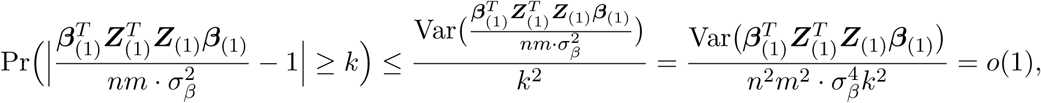

and

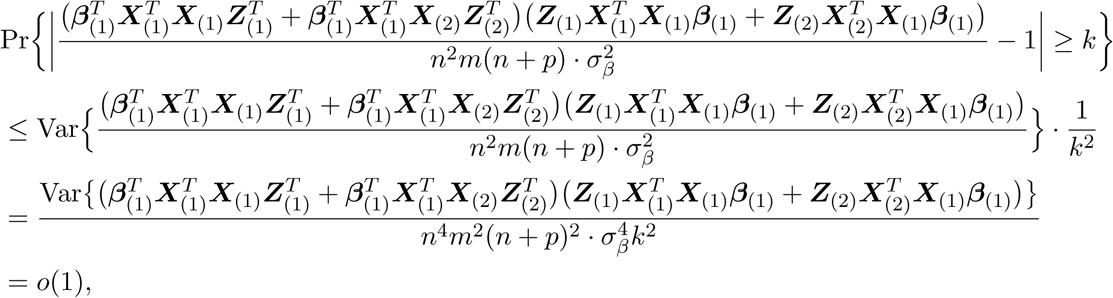

and

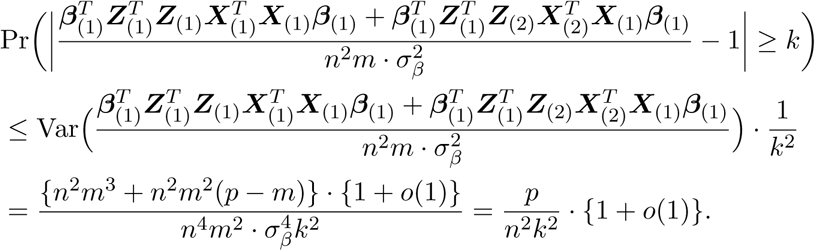

In follows that Proposition A1 is proved. More generally, if training and testing data have different sample sizes, donated as *n* and *n*_*z*_, respectively, we have the following results.

### Proposition S2.

*Under Condition (1), if m* → *∞ as n, n*_*z*_, *p* → *∞, then we have*

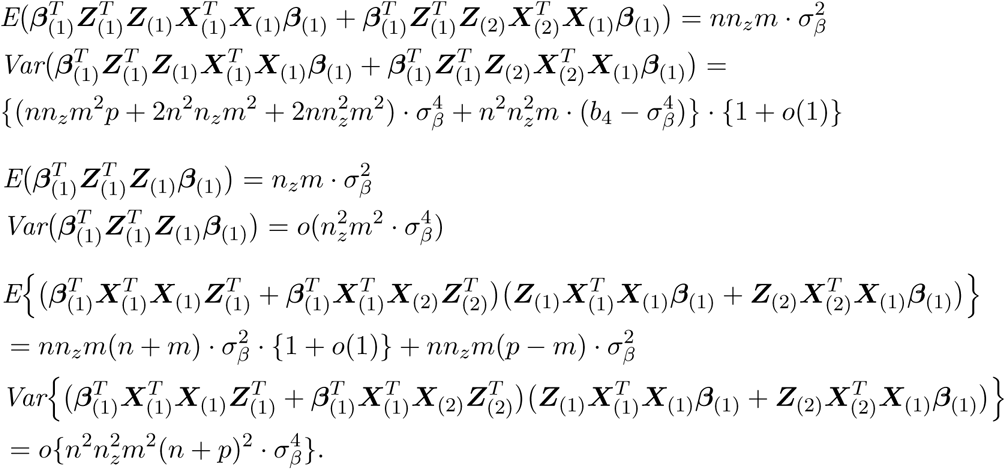

By Markov’s inequality and continuous mapping theorem again, we have the following results.

### Proposition S3.

*Under Condition (1), if m* → *∞ as n, n*_*z*_, *p* → *∞, then we have*

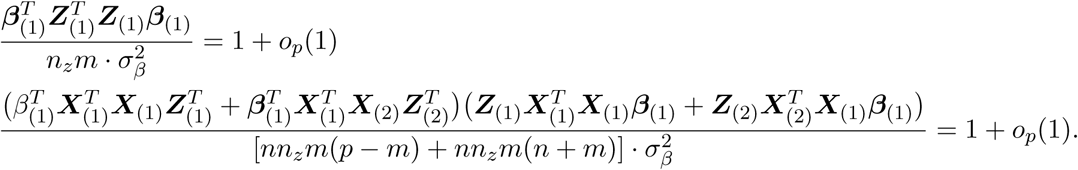

*If further p/*(*nn*_*z*_) → 0, *then we have*

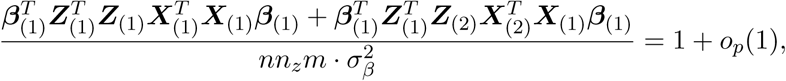

*and it follows that*

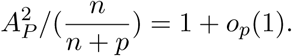

When *p/*(*nn*_*z*_) ↛ 0, i.e., *α* ∈ [1, *∞*], we note that

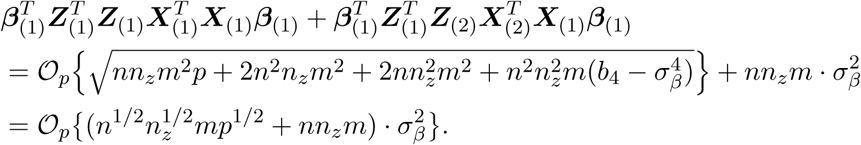

It follows that

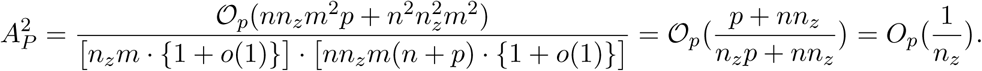

The results of threshold-PRS can also be derived in a similar way. Without loss of generality, we set 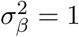 in later steps.

### Proposition S4.

*Under Conditions (1), if m, q*_1_, *q*_2_ → *∞ as n, p* → *∞, then we have*

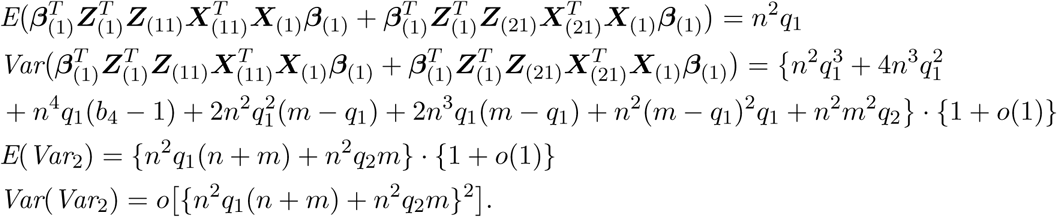

By Markov’s inequality and continuous mapping theorem again, Proposition A2 is proved.

## 2. Technical details

The following technical details are useful to prove our theoretical results. Most of them involve calculating the asymptotic expectation of the trace of the product of multiple large random matrices. We use the definition of matrix trace and apply the combination theory to calculate the total variations. The results provided below may also benefit other research questions involving similar calculations.

### 2.1 GWAS-PRS

First moment of covariance term

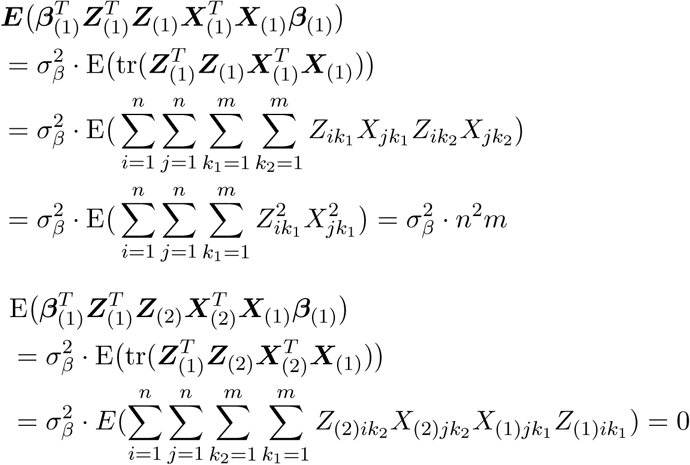

Thus, we have 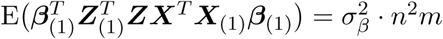.

#### First moment of variance term I

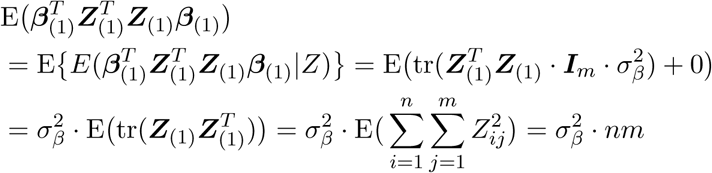

#### First moment of variance term II

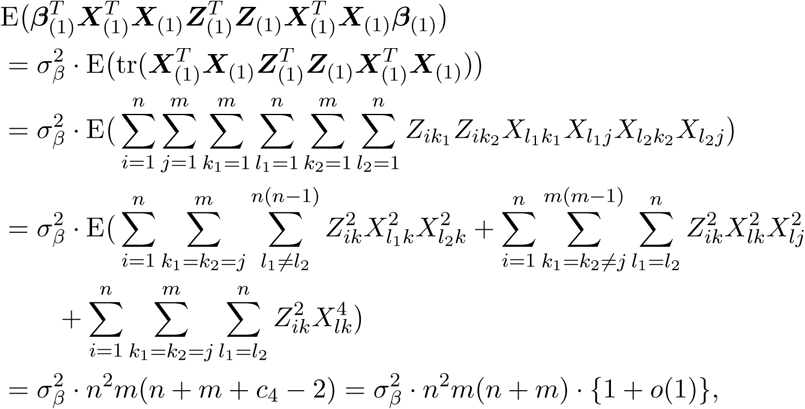

where 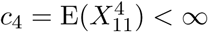

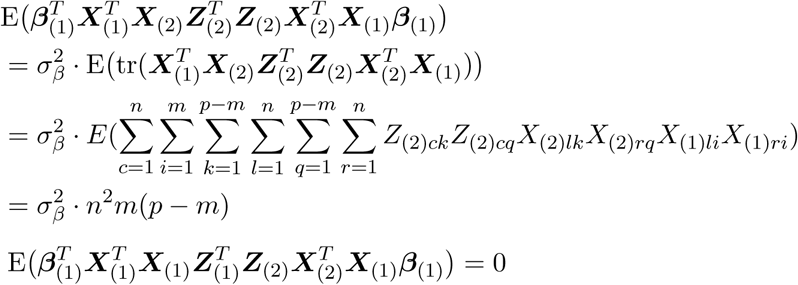

Thus, we have

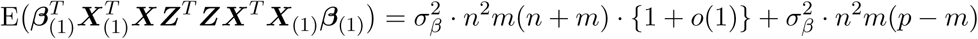

#### Second moment of covariance term

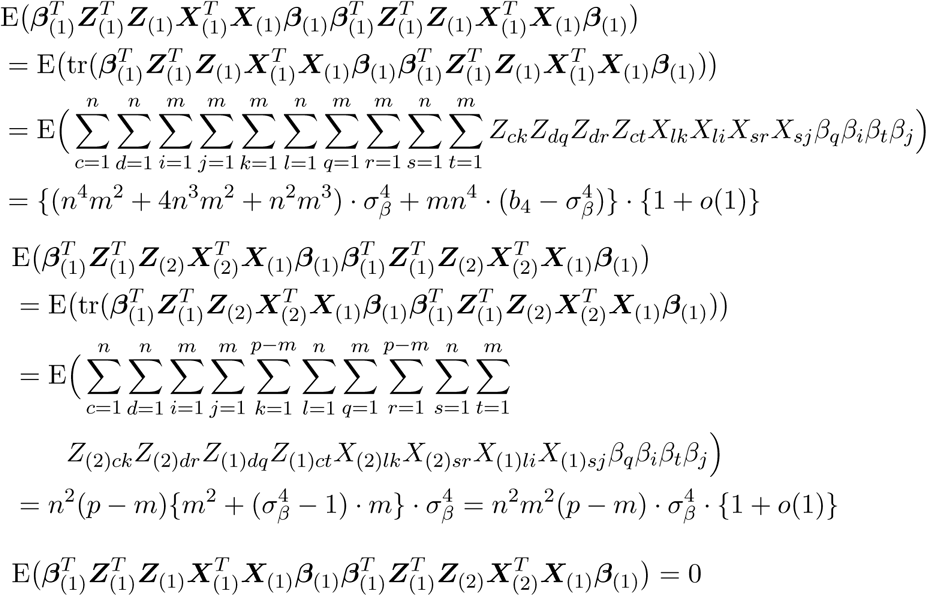

It follows that

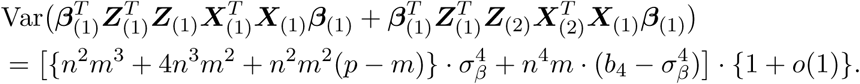

#### Second moment of variance term I

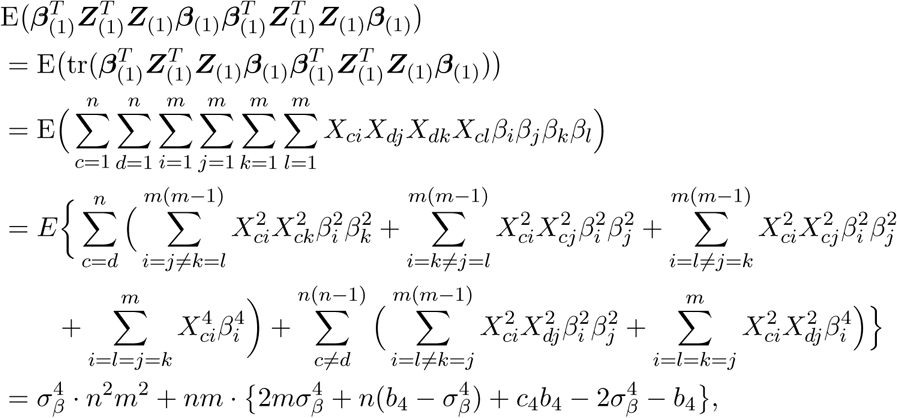

Where 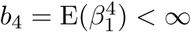. It follows that

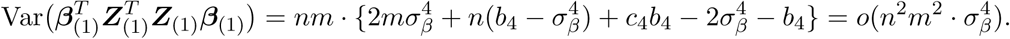

#### Second moment of variance term II

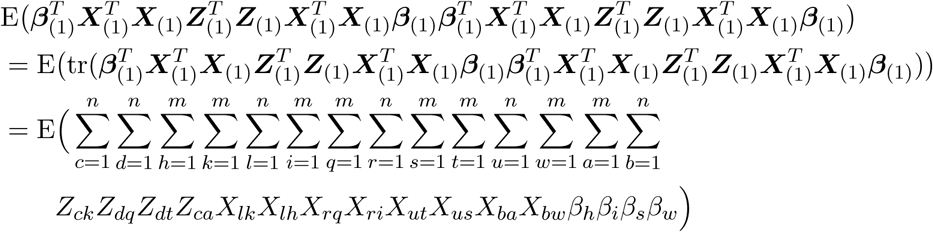

To have non-zero means, we need possible combinations with the following patterns:

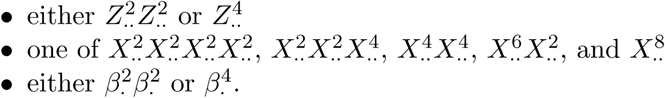

After tedious calculations, we have

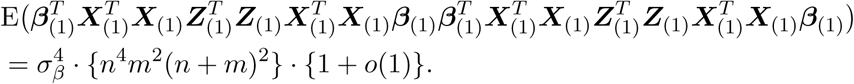

Next, we have

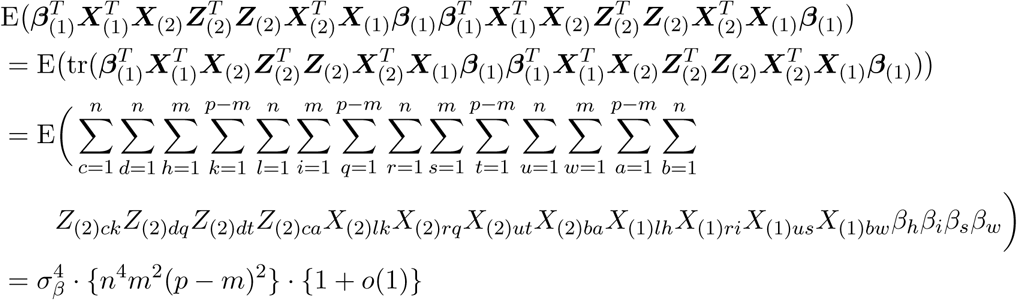

and

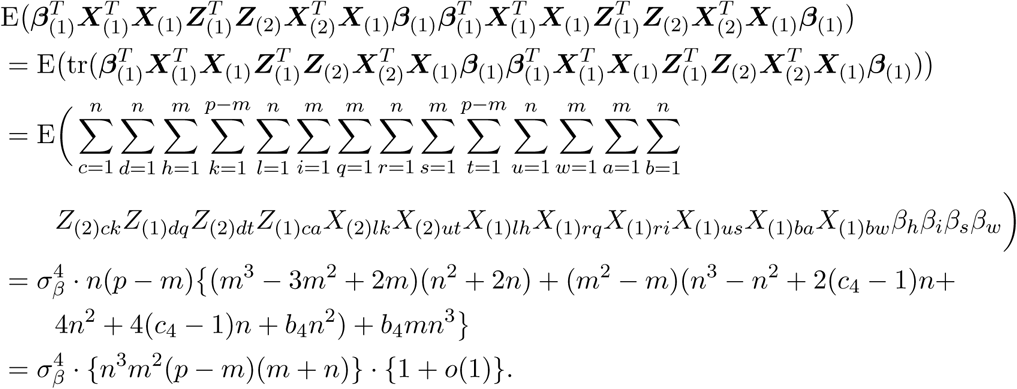

Similarly, we have

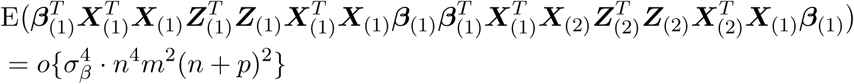

and

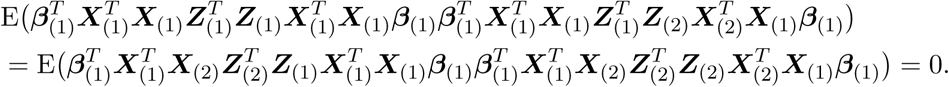

It follows that

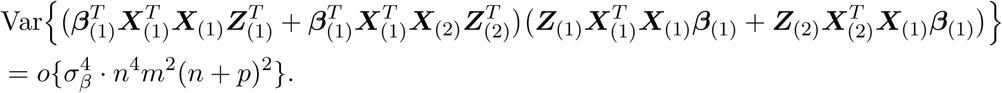

The results of different sample sizes can be similarly derived and are ignored. Without loss of generality, we set 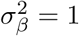 for simplicity below for threshold-PRS.

### 2.2 Threshold-PRS

**First moment of covariance term**

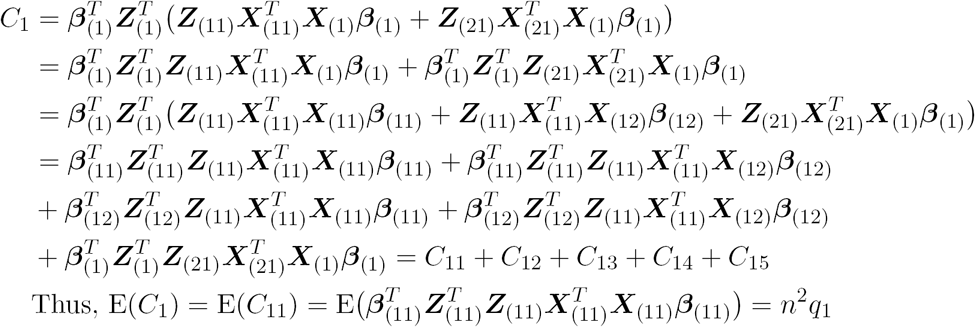

#### First moment of variance term II

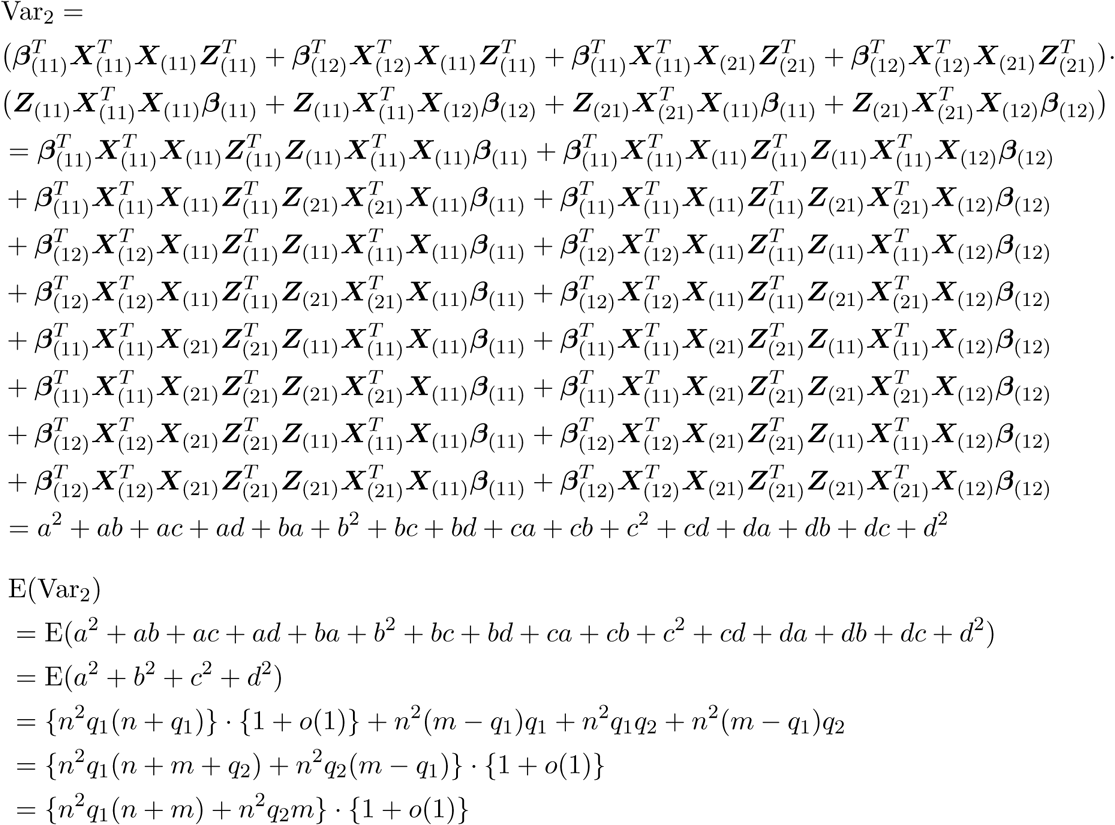

#### Second moment of covariance term

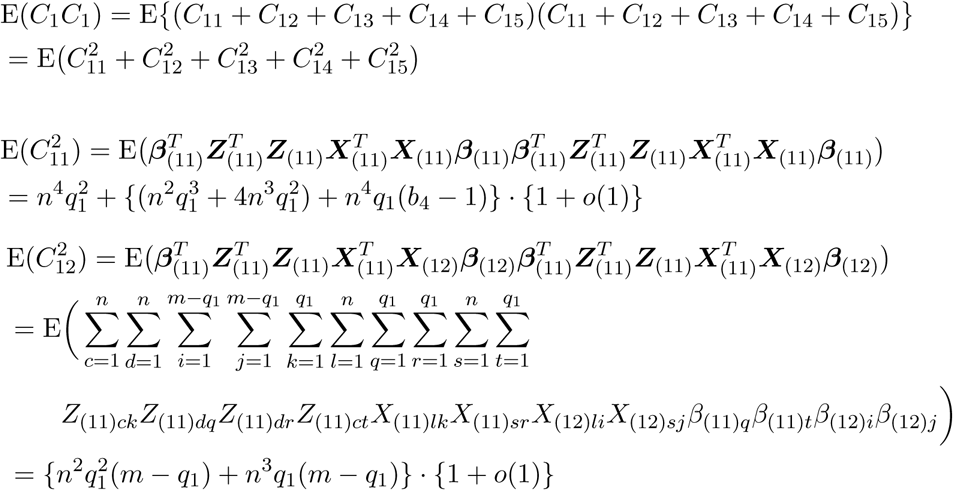

Similarly, we have

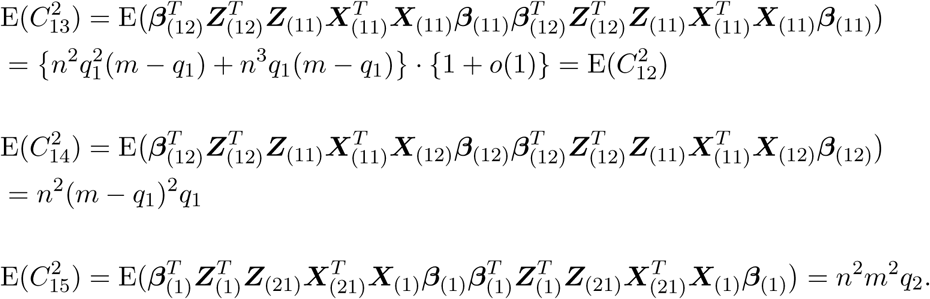

Thus, we have

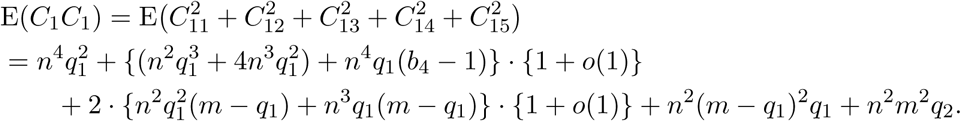

It follows that

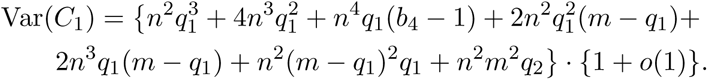

#### Second moment of variance term II

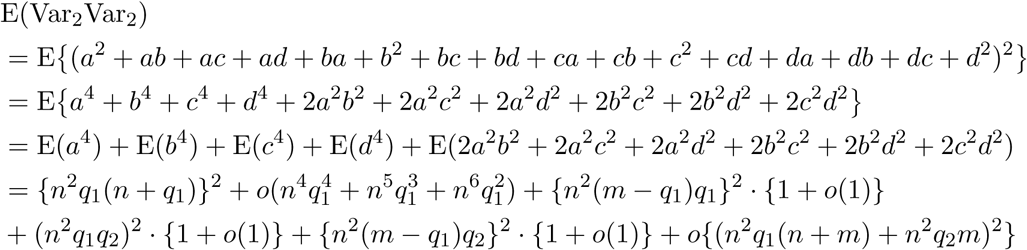

It follows that Var(var_2_)=*o*[{*n*^2^*q*1(*n*+*m*) + *n*^2^*q*2*m*}^2^].

## 3. Simulation setups: BLUP, Ridge, and LASSO

We perform simple simulations to numerically illustrate the prediction accuracy of BLUP, ridge and LASSO given different *m/p* ratios.

As in Case 1, we simulate *p* = 100, 000 uncorrelated SNPs from *N* (0, 1). We vary the number of causal SNPs *m* and set it to 100, 1000, 10, 000 and 50, 000. The casual SNP effects ***β***_(1)_ ∼ *MV N* (0, ***I***_*m*_). The linear polygenic model (2) is used to generate phenotype ***y*.** The sample size is set to 1000 or 10, 000 for training data, and 1000 for testing data. A total of 100 replications are conducted for each simulation condition. The results of BLUP prediction are shown in Supplementary Figure 18. The performance of BLUP is consistent across different *m/p* ratios, and is slightly larger than 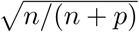

For LASSO and ridge regression, we use smaller (*n, p, m*) to reduce the computational burden. We simulate *p* = 10, 000 uncorrelated SNPs from *N* (0, 1). We vary the number of causal SNPs *m* and set it to 100, 1000, 25, 000 and 5, 000. The sample size is set to 2000 for training data and 1000 for testing data. Other settings are exactly the same as those in BLUP. LASSO has nearly perfect performance when the signals are sparse. However, its performance is consistently lower than 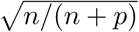 when the signals are dense (Supplementary Figure 19). The pattern of ridge regression is very similar to that of PRS. When all the SNPs are used, ridge regression has consistent perfomrance regardless of the *m/p* ratios, which is slightly larger than 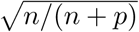 (Supplementary Figure 20).

## 4. Real data analysis

The detailed cohort information, data quality control, and GWAS procedures can be found in Zhao et al. (2019), Supplementary Table 1 gives a brief summary of the demographic information of five datasets. In each testing dataset, we use PLINK toolset (Purcell et al., 2007) to generate risk scores in testing data by summarizing across pruned (window size 50, step 5, *R*^2^ = 0.2) SNP alleles, weighed by their effect sizes estimated from training data. The UKB effect size estimates are taken from Zhao et al. (2019) and have been made publicly available at https://med.sites.unc.edu/bigs2/data/gwas-summary-statistics/. The association between each pair of polygenic profile and ROI volume is estimated and tested in linear regression, adjusting for the age and gender. The additional variance of ROI volume that can be explained by PRS is used to measure the prediction power.

### Data Acknowledgement

Part of data collection and sharing for this project was funded by the Alzheimers Disease Neuroimaging initiative (ADNI) (National Institutes of Health Grant U01 AG024904) and DOD ADNI (Department of Defense award number W81XWH-12-2-0012). ADNI is funded by the National Institute on Aging, the National Institute of Biomedical Imaging and Bioengineering and through generous contributions from the following: Alzheimers Association; Alzheimers Drug Discovery Foundation; Araclon Biotech; BioClinica, Inc.; Biogen Idec Inc.; Bristol-Myers Squibb Company; Eisai Inc.; Elan Pharmaceuticals, Inc.; Eli Lilly and Company; EuroImmun; F. Hoffmann-La Roche Ltd and its affiliated company Genentech, Inc.; Fujirebio; GE Healthcare; IXICO Ltd; Janssen Alzheimer Immunotherapy Research & Development, LLC; Johnson & Johnson Pharmaceutical Research & Development LLC; Medpace, Inc.; Merck & Co., Inc.; Meso Scale Diagnostics, LLC; NeuroRx Research; Neurotrack Technologies; Novartis Pharmaceuticals Corporation; Pfizer Inc.; Piramal Imaging; Servier; Synarc Inc.; and Takeda Pharmaceutical Company. The Canadian Institutes of Health Research is providing funds to support ADNI clinical sites in Canada. Private sector contributions are facilitated by the Foundation for the National Institutes of Health. The grantee organization is the Northern California Institute for Research and Education, and the study is coordinated by the Alzheimers Disease Cooperative Study at the University of California, San Diego. ADNI data are disseminated by the Laboratory for Neuro Imaging at the University of Southern California. Part of the data collection and sharing for this project was funded by the Pediatric Imaging, Neurocognition and Genetics Study (PING) (U.S. National Institutes of Health Grant RC2DA029475). PING is funded by the National Institute on Drug Abuse and the Eunice Kennedy Shriver National Institute of Child Health & Human Development. PING data are disseminated by the PING Coordinating Center at the Center for Human Development, University of California, San Diego. HCP data were provided by the Human Connectome Project, WU-Minn Consortium (Principal Investigators: David Van Essen and Kamil Ugurbil; 1U54MH091657) funded by the 16 NIH Institutes and Centers that support the NIH Blueprint for Neuroscience Research; and by the McDonnell Center for Systems Neuroscience at Washington University.

**Supplementary Table 1.** Demographic information of five datasets.

| Dataset | Sample size | Mean age (s.d.) | Age range | Proportion of male |
| --- | --- | --- | --- | --- |
| UK Biobank | 19,629 | 62.51 (7.47) | (40 80) | 0.473 |
| ADNI | 1248 | 74.18 (7.16) | (55 92) | 0.566 |
| HCP | 1141 | 28.82 (3.68) | (22 36) | 0.542 |
| PING | 924 | 12.28 (4.99) | (3 21) | 0.518 |
| PNC | 1492 | 21.14 (3.71) | (14 29) | 0.515 |

### PING Methods

Part of the data used in the preparation of this article were obtained from the Pediatric Imaging, Neurocognition and Genetics (PING) Study database (http://ping.chd.ucsd.edu/). PING was launched in 2009 by the National Institute on Drug Abuse (NIDA) and the Eunice Kennedy Shriver National Institute Of Child Health & Human Development (NICHD) as a 2-year project of the American Recovery and Reinvestment Act. The primary goal of PING has been to create a data resource of highly standardized and carefully curated magnetic resonance imaging (MRI) data, comprehensive genotyping data, and developmental and neuropsychological assessments for a large cohort of developing children aged 3 to 20 years. The scientific aim of the project is, by openly sharing these data, to amplify the power and productivity of investigations of healthy and disordered development in children, and to increase understanding of the origins of variation in neurobehavioral phenotypes. For up-to-date information, see http://ping.chd.ucsd.edu/.

### ADNI Methods

Data used in the preparation of this article were obtained from the Alzheimers Disease Neuroimaging Initiative (ADNI) database (http://adni.loni.usc.edu). The ADNI was launched in 2003 by the National Institute on Aging (NIA), the National Institute of Biomedical Imaging and Bioengineering (NIBIB), the Food and Drug Administration (FDA), private pharmaceutical companies and non-profit organizations, as a 60 million, 5-year public-private partnership. The primary goal of ADNI has been to test whether serial magnetic resonance imaging (MRI), positron emission tomography (PET), other biological markers, and clinical and neuropsychological assessment can be combined to measure the progression of mild cognitive impairment (MCI) and early Alzheimers disease (AD). Determination of sensitive and specific markers of very early AD progression is intended to aid researchers and clinicians to develop new treatments and monitor their effectiveness, as well as lessen the time and cost of clinical trials.

The Principal Investigator of this initiative is Michael W. Weiner, MD, VA Medical Center and University of California San Francisco. ADNI is the result of efforts of many co-investigators from a broad range of academic institutions and private corporations, and subjects have been recruited from over 50 sites across the U.S. and Canada. The initial goal of ADNI was to recruit 800 subjects but ADNI has been followed by ADNI-GO and ADNI-2. To date these three protocols have recruited over 1500 adults, ages 55 to 90, to participate in the research, consisting of cognitively normal older individuals, people with early or late MCI, and people with early AD. The follow up duration of each group is specified in the protocols for ADNI-1, ADNI-2 and ADNI-GO. Subjects originally recruited for ADNI-1 and ADNI-GO had the option to be followed in ADNI-2. For up-to-date information, see www.adni-info.org.

*Pediatric Imaging, Neurocognition and Genetics (PING) Authors*. Connor McCabe^1^, Linda Chang^2^, Natacha Akshoomoff^3^, Erik Newman^1^, Thomas Ernst^2^, Peter Van Zijl^4^, Joshua Kuperman^5^, Sarah Murray^6^, Cinnamon Bloss^6^, Mark Appelbaum^1^, Anthony Gamst^1^, Wesley Thompson^3^, Hauke Bartsch^5^.

*Alzheimer’s Disease Neuroimaging Initiative (ADNI) Authors*. Michael Weiner^7^, Paul Aisen^1^, Ronald Petersen^8^, Clifford R. Jack Jr^8^, William Jagust^9^, John Q. Trojanowki^10^, Arthur W. Toga^11^, Laurel Beckett^12^, Robert C. Green^13^, Andrew J. Saykin^14^, John Morris^15^, Leslie M. Shaw^10^, Zaven Khachaturian^16^, Greg Sorensen^17^, Maria Carrillo^18^, Lew Kuller^19^, Marc Raichle^15^, Steven Paul^20^, Peter Davies^21^, Howard Fillit^22^, Franz Hefti^23^, Davie Holtzman^15^, M. Marcel Mesulman^24^, William Potter^25^, Peter J. Snyder^26^, Adam Schwartz^27^, Tom Montine^28^, Ronald G. Thomas^1^, Michael Donohue^1^, Sarah Walter^1^, Devon Gessert^1^, Tamie Sather^1^, Gus Jiminez^1^, Danielle Harvey^12^, Matthew Bernstein^8^, Nick Fox^29^, Paul Thompson^11^, Norbert Schuff^7^, Charles DeCarli^12^, Bret Borowski^8^, Jeff Gunter^8^, Matt Senjem^8^, Prashanthi Vemuri^8^, David Jones^8^, Kejal Kantarci^8^, Chad Ward^8^, Robert A. Koeppe^30^, Norm Foster^31^, Eric M. Reiman^32^, Kewei Chen^32^, Chet Mathis^19^, Susan Landau^9^, Nigel J. Cairns^15^, Erin Householder^15^, Lisa Taylor-Reinwald^15^, Virginia M.Y. Lee^10^, Magdalena Korecka^10^, Michal Figurski^10^, Karen Crawford^11^, Scott Neu^11^, Tatiana M. Foroud^14^, Steven Potkin^33^, Li Shen^14^, Kelley Faber^14^, Sungeun Kim^14^, Kwangsik Nho^14^, Leon Thal^1^, Richard Frank^34^, Neil Buckholtz^35^, Marilyn Albert^36^, John Hsiao^35^.

^1^UC San Diego, La Jolla, CA 92093, USA. ^2^U Hawaii, Honolulu, HI 96822, USA. ^3^Department of Psychiatry, University of California, San Diego, La Jolla, California 92093, USA. ^4^Kennedy Krieger Institute, Baltimore, MD 21205, USA. ^5^Multimodal Imaging Laboratory, Department of Radiology, University of California San Diego, La Jolla, California 92037, USA. ^6^Scripps Translational Science Institute, La Jolla, CA 92037, USA. ^7^UC San Francisco, San Francisco, CA 94143, USA. ^8^Mayo Clinic, Rochester, MN 55905, USA. ^9^UC Berkeley, Berkeley, CA 94720-5800, USA. ^10^U Pennsylvania, Philadelphia, PA 19104, USA. ^11^USC, University of Southern California, Los Angeles, CA 90033, USA. ^12^UC Davis, Davis, CA 95616, USA. ^13^Brigham and Women s Hospital/Harvard Medical School, Boston MA 02115, USA. ^14^Indiana University, Indianapolis, IN 46202-5143, USA. ^15^Washington University St. Louis, St. Louis, MO 63130, USA. ^16^Prevent Alzheimers Disease 2020, Rockville, MD 20850, USA. ^17^Siemens ^18^Alzheimers Association, Chicago, IL 60601, USA. ^19^University of Pittsburgh, Pittsburgh, PA 15260, USA. ^20^Cornell University, Ithaca, NY 14850, USA. ^21^Albert Einstein College of Medicine of Yeshiva University, Bronx, NY 10461, USA. ^22^AD Drug Discovery Foundation, New York, NY 10019, USA. ^23^Acumen Pharmaceuticals, Livermore, California 94551, USA. ^24^Northwestern University, Evanston, IL 60208, USA. ^25^National Institute of Mental Health, Bethesda, MD 20892-9663, USA. ^26^Brown University, Providence, RI 02912, USA. ^27^Eli Lilly, Indianapolis, Indiana 46285, USA. ^28^University of Washington, Seattle, WA 98195, USA. ^29^University of London, London WC^1^E ^7^HU, UK. ^30^University of Michigan, Ann Arbor, MI 48109, USA. ^31^University of Utah, Salt Lake City, UT 84112, USA. ^32^Banner Alzheimers Institute, Phoenix, AZ 85006, USA. ^33^UC Irvine, Irvine, CA 92697, USA. ^34^General Electric ^35^National Institute on Aging/National Institutes of Health, Bethesda, MD 20892, USA. ^36^The Johns Hopkins University, Baltimore, MD 21218, USA.

---

**Supplementary Figure 1:**
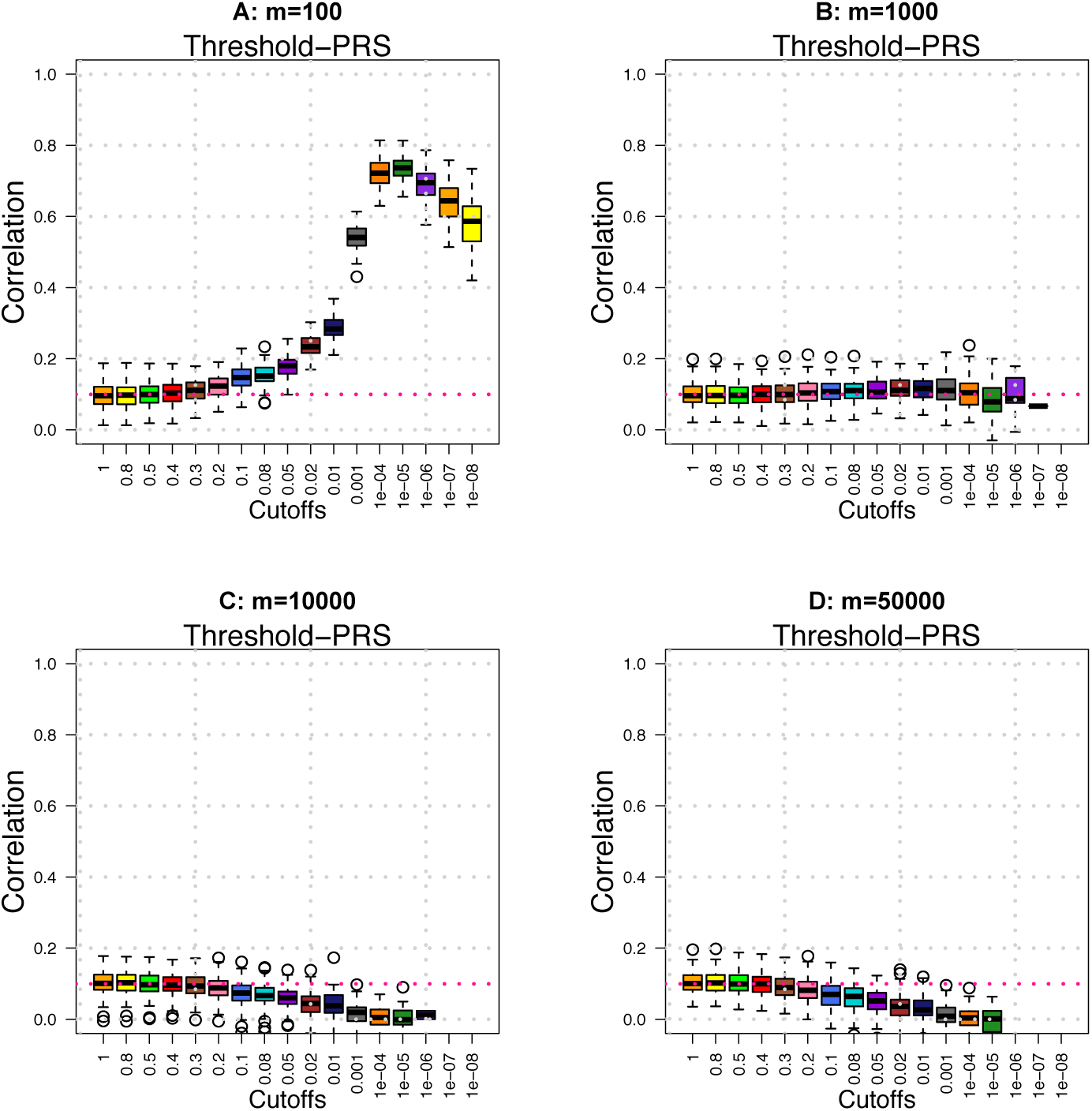
Prediction accuracy (*A*_*P*_) of threshold-PRS across different *m/p* ratios. We set *p*=100, 000 and *n*=1000 in both training and testing data.

**Supplementary Figure 2:**
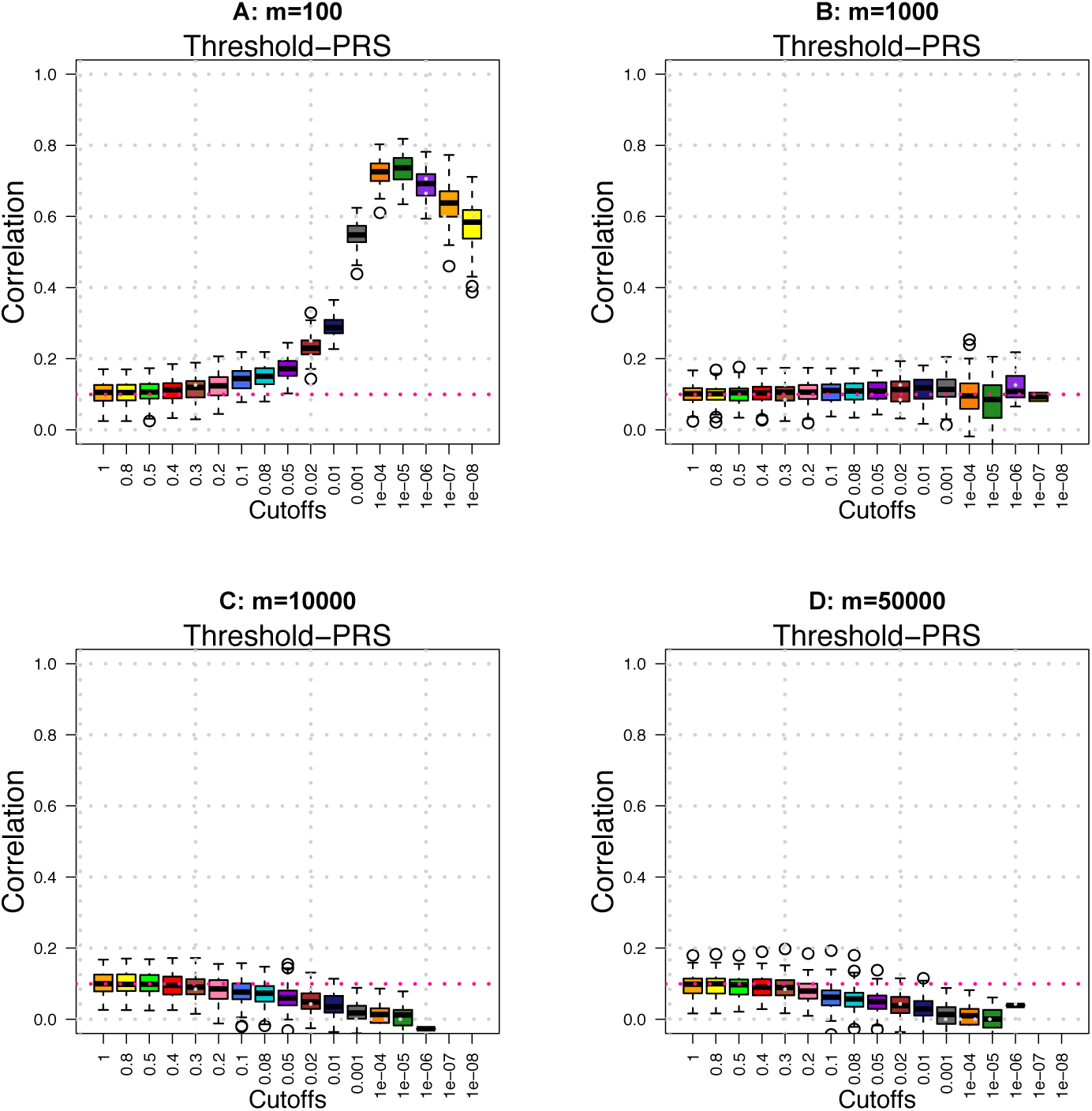
Prediction accuracy (*A*_*P*_) of threshold-PRS across different *m/p* ratios. SNP data are independently sampled from {0, 1, 2}. We set *p*=100, 000 and *n*=1000 in both training and testing data.

**Supplementary Figure 3:**
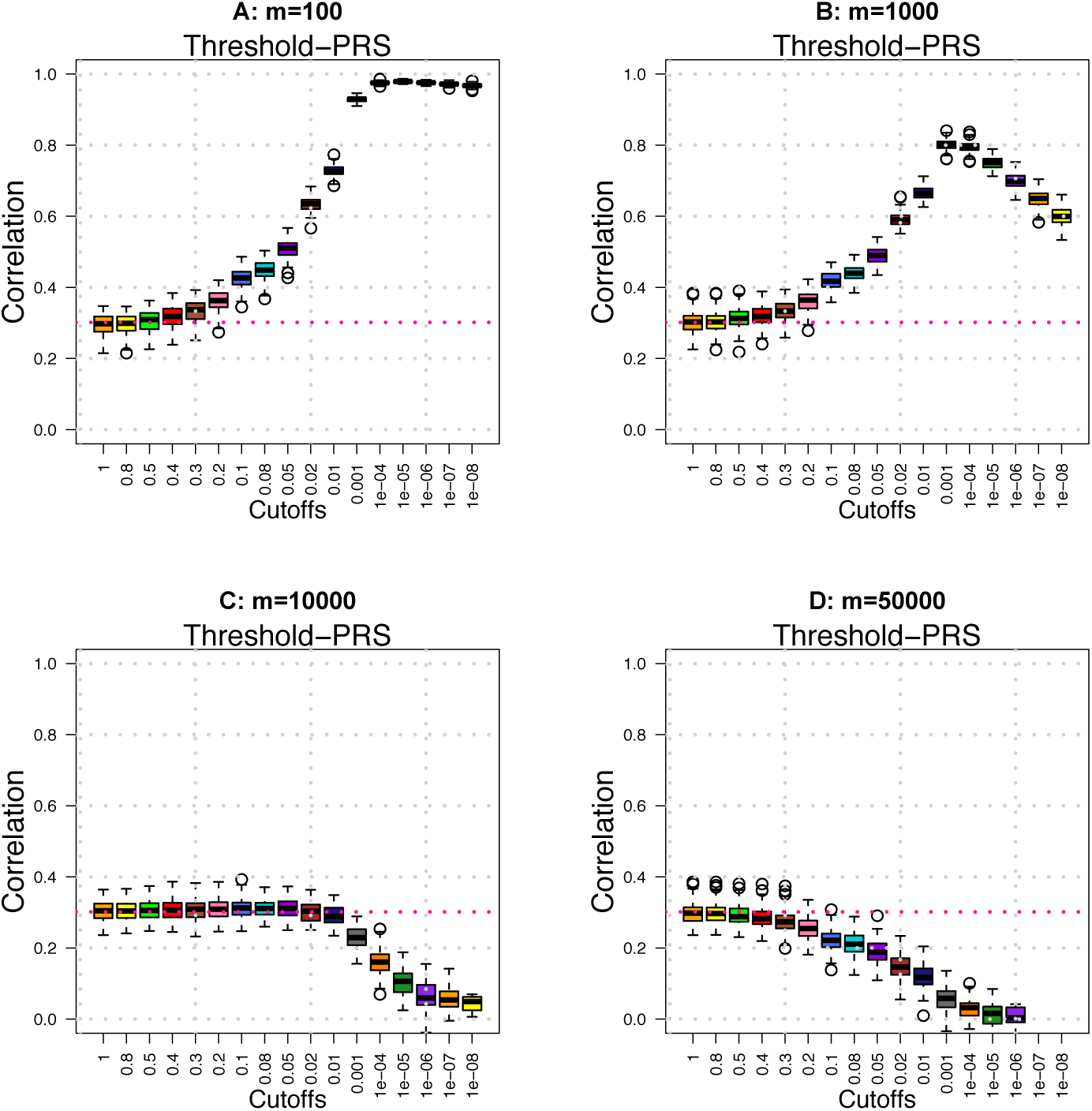
Prediction accuracy (*A*_*P*_) of threshold-PRS across different *m/p* ratios. SNP data are independently sampled from {0, 1, 2}. We set *p*=100, 000 and *n*=10, 000 in training data and *n* = 1000 in testing data.

**Supplementary Figure 4:**
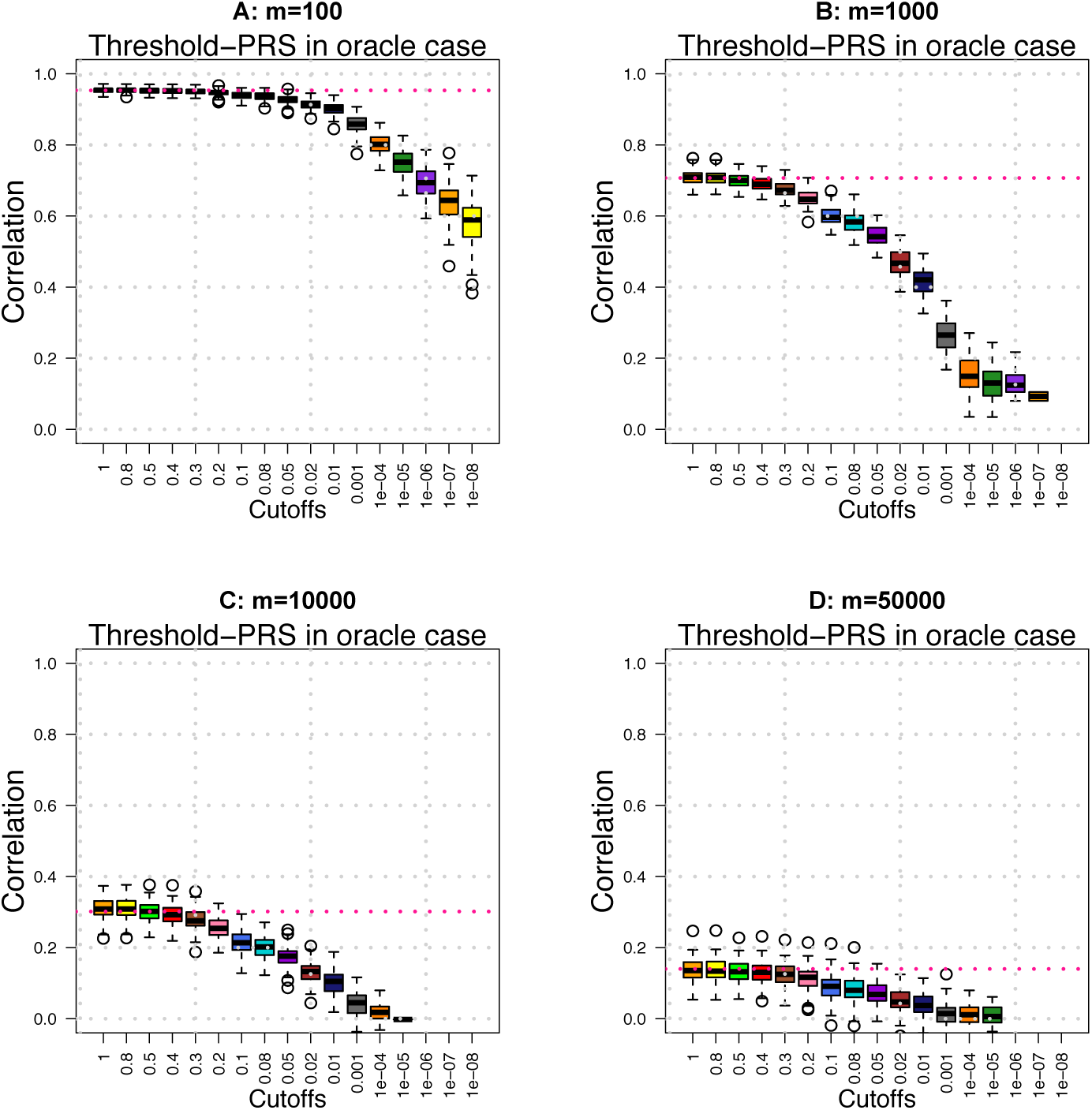
Prediction accuracy (*A*_*P*_) of threshold-PRS across different *m/p* ratios when the *m* causal SNPs are known and are only considered as candidates in constructing PRS. SNP data are independently sampled from {0, 1, 2}. We set *p*=100, 000 and *n*=1000 in both training and testing data.

**Supplementary Figure 5:**
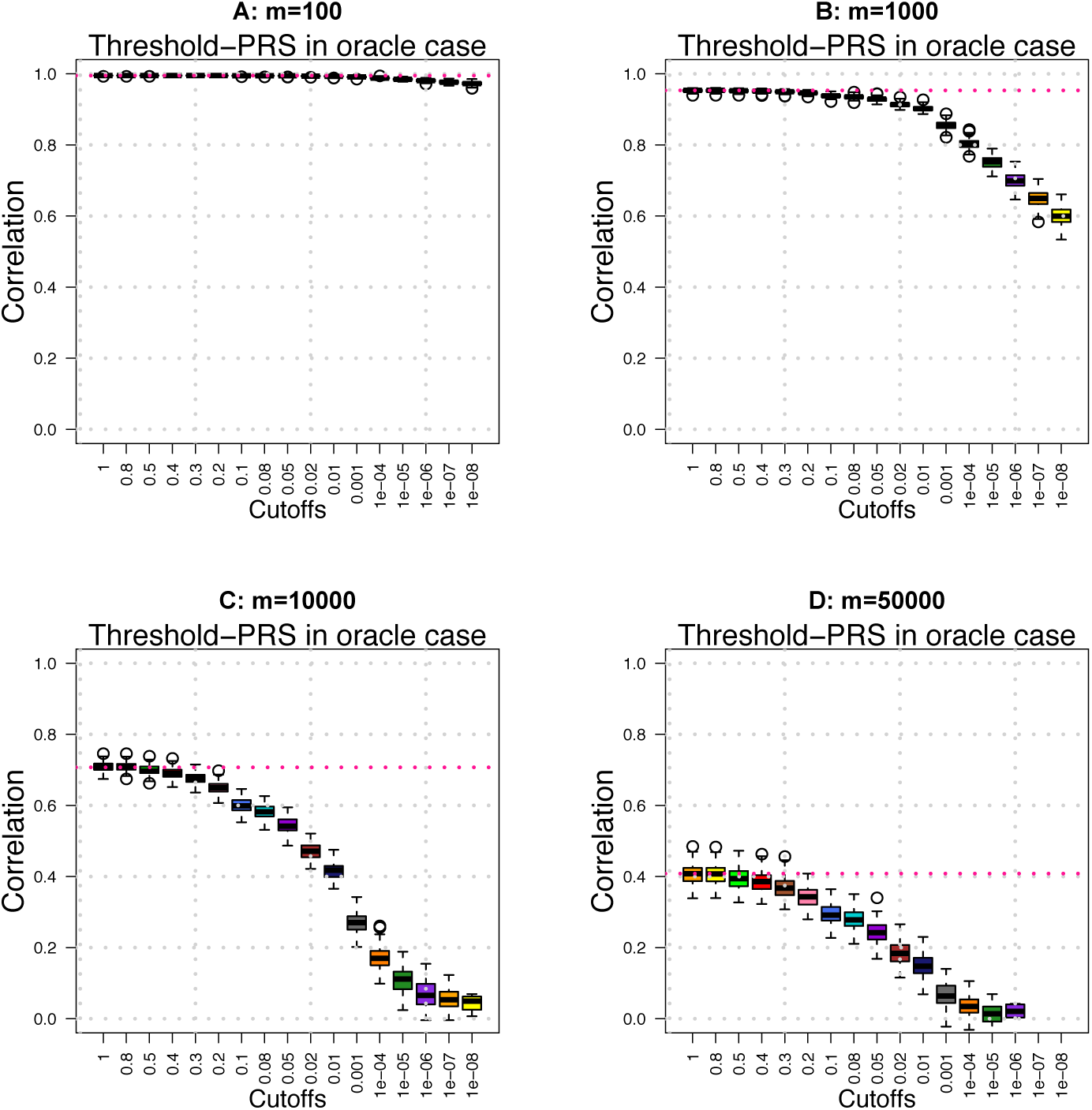
Prediction accuracy (*A*_*P*_) of threshold-PRS across different *m/p* ratios when the *m* causal SNPs are known and are only considered as candidates in constructing PRS. SNP data are independently sampled from {0, 1, 2}. We set *p*=100, 000 and *n*=10, 000 in training data and *n* = 1000 in testing data.

**Supplementary Figure 6:**
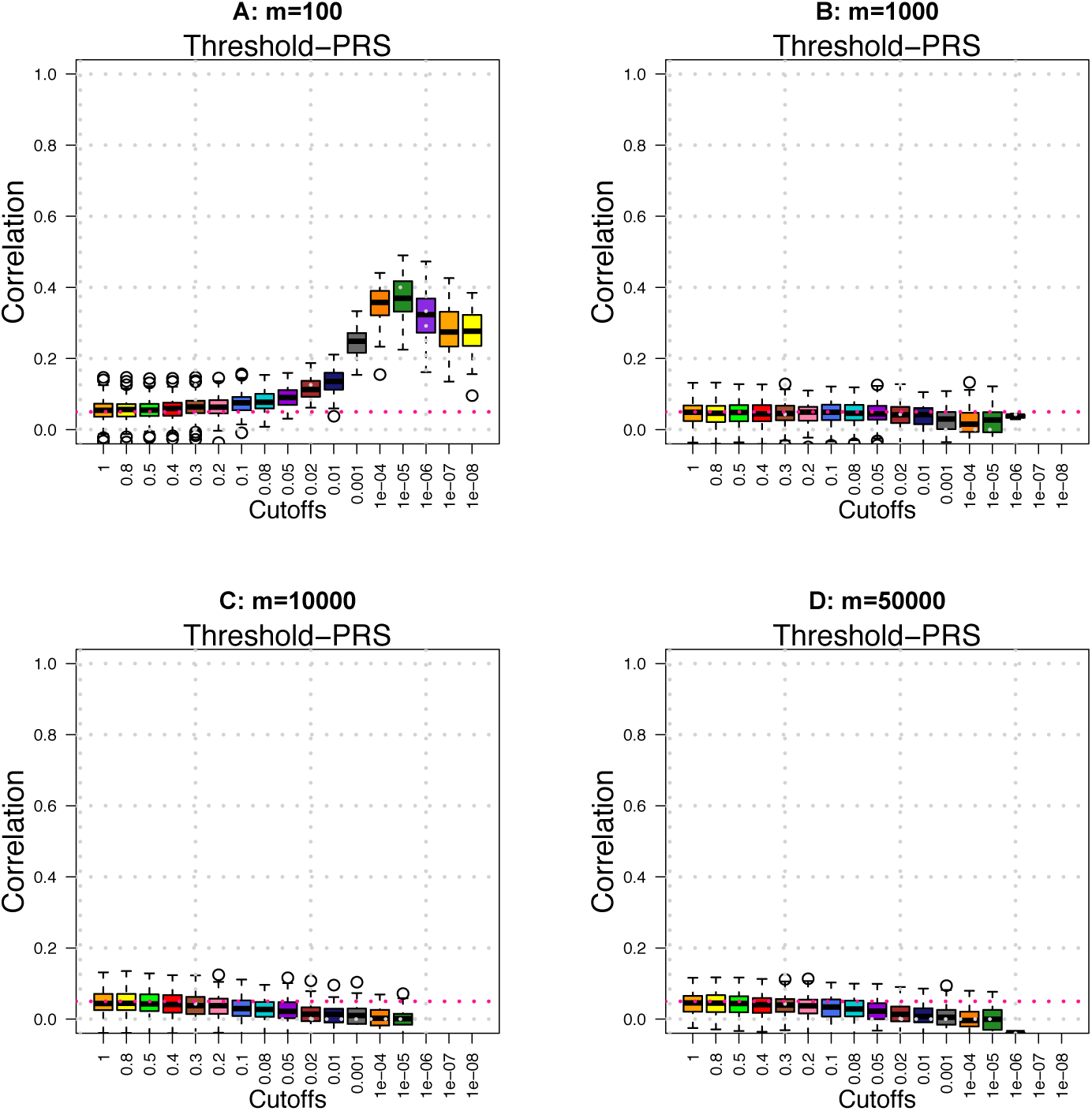
Prediction accuracy (*A*_*P*_) of threshold-PRS across different *m/p* ratios when the heritability *h*^2^=0.5. SNP data are independently sampled from {0, 1, 2}. We set *p*=100, 000 and *n*=1000 in both training and testing data.

**Supplementary Figure 7:**
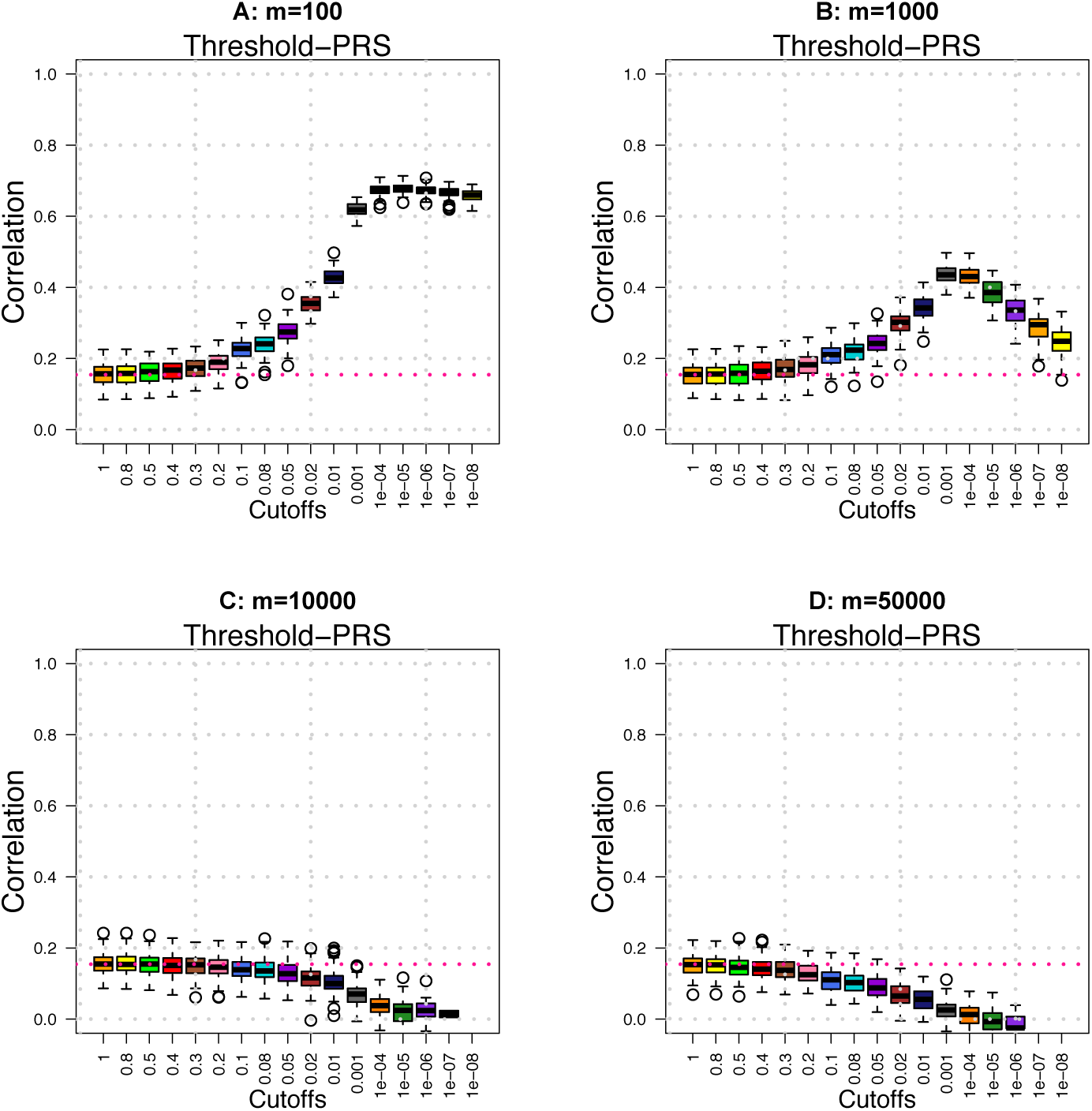
Prediction accuracy (*A*_*P*_) of threshold-PRS across different *m/p* when the heritability *h*^2^=0.5. SNP data are independently sampled from {0, 1, 2}. We set *p*=100, 000 and *n*=10, 000 in training data and *n* = 1000 in testing data.

**Supplementary Figure 8:**
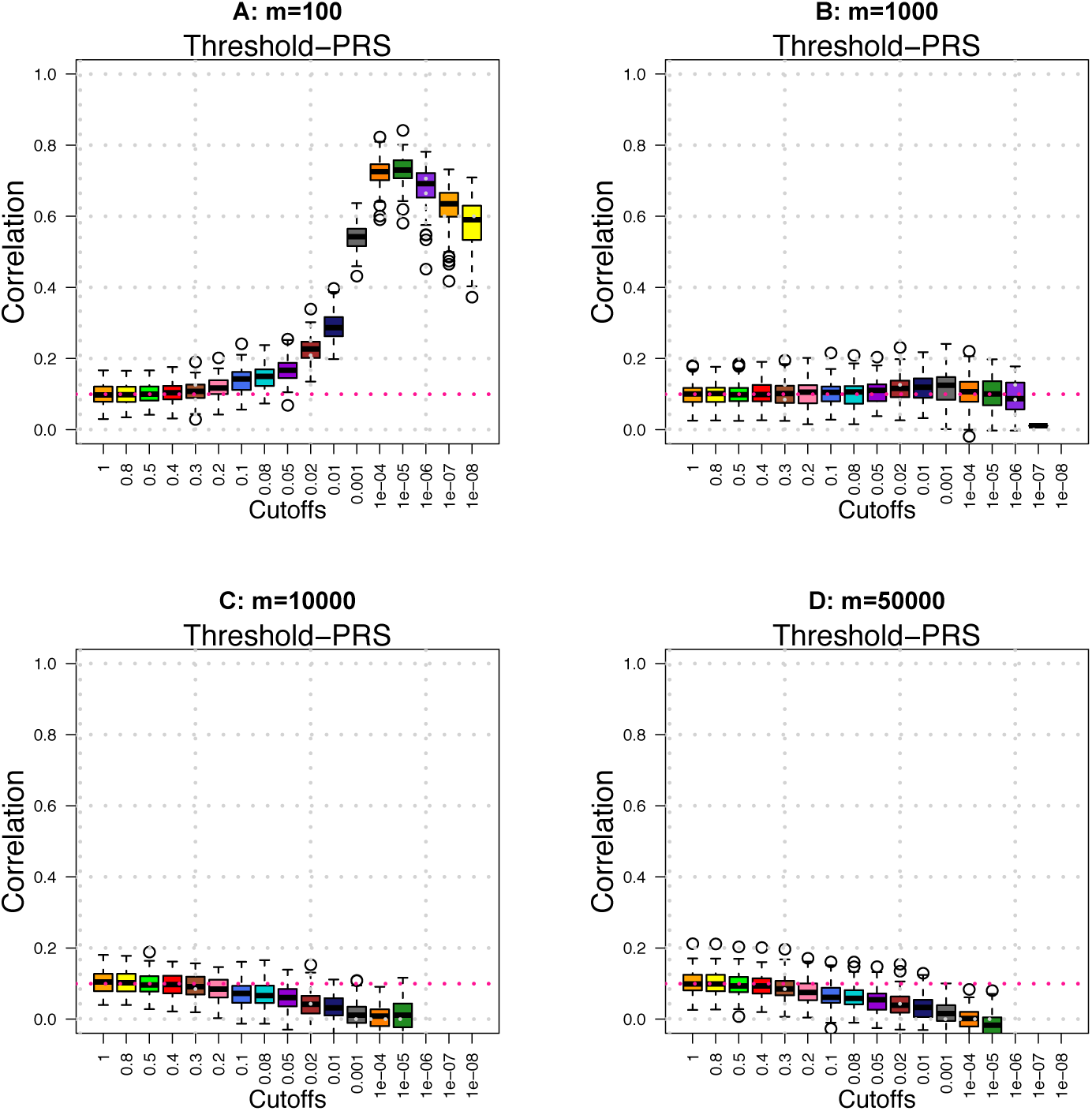
Prediction accuracy (*A*_*P*_) of threshold-PRS across different *m/p* ratios when population substructure exists in the SNP data. SNP data are independently sampled from {0, 1, 2}. We set *p*=100, 000 and *n*=1000 in both training and testing data.

**Supplementary Figure 9:**
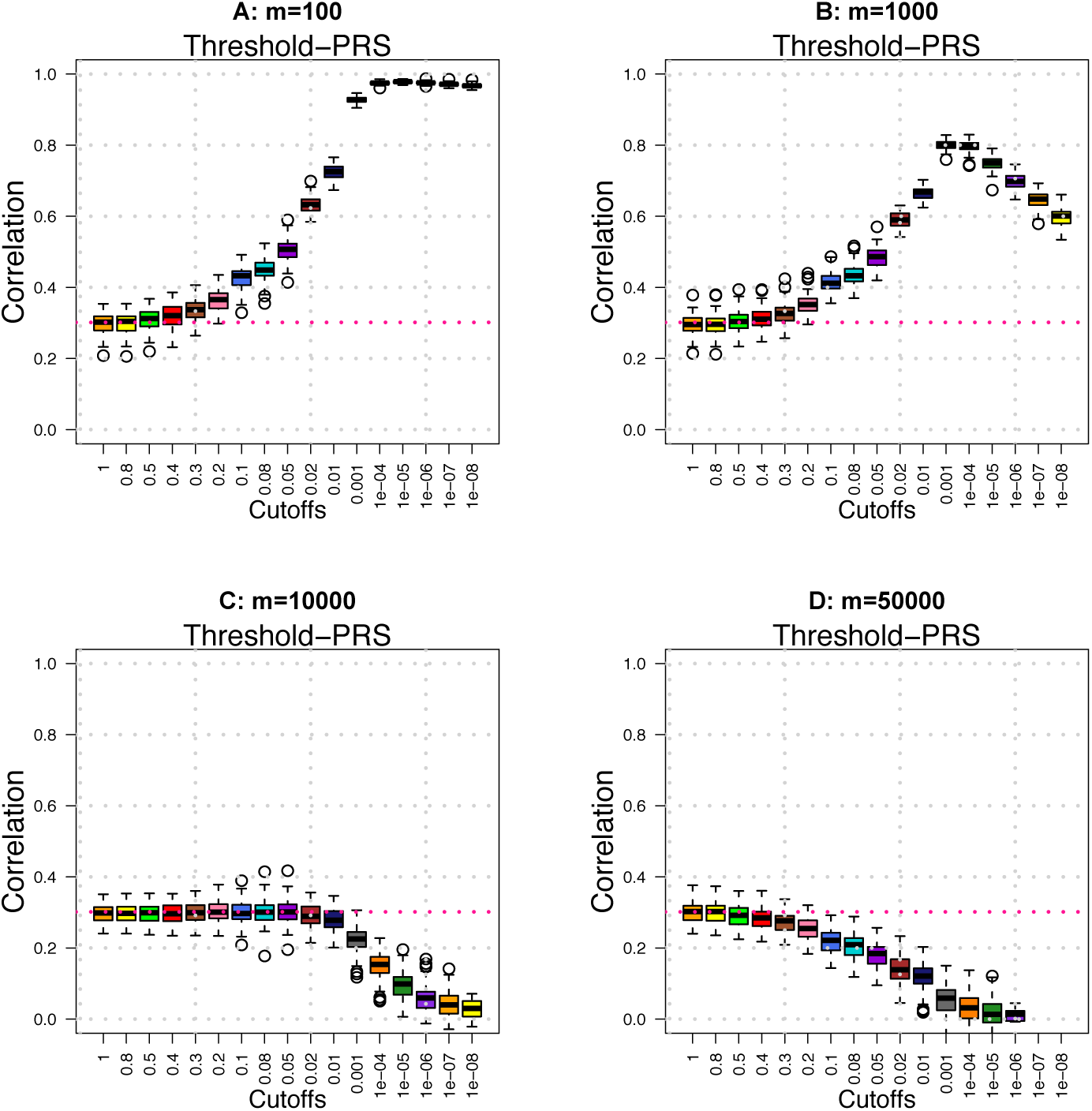
Prediction accuracy (*A*_*P*_) of threshold-PRS across different *m/p* ratios when population substructure exists in the SNP data. SNP data are independently sampled from {0, 1, 2}. We set *p*=100, 000 and *n*=10, 000 in training data and *n* = 1000 in testing data.

**Supplementary Figure 10:**
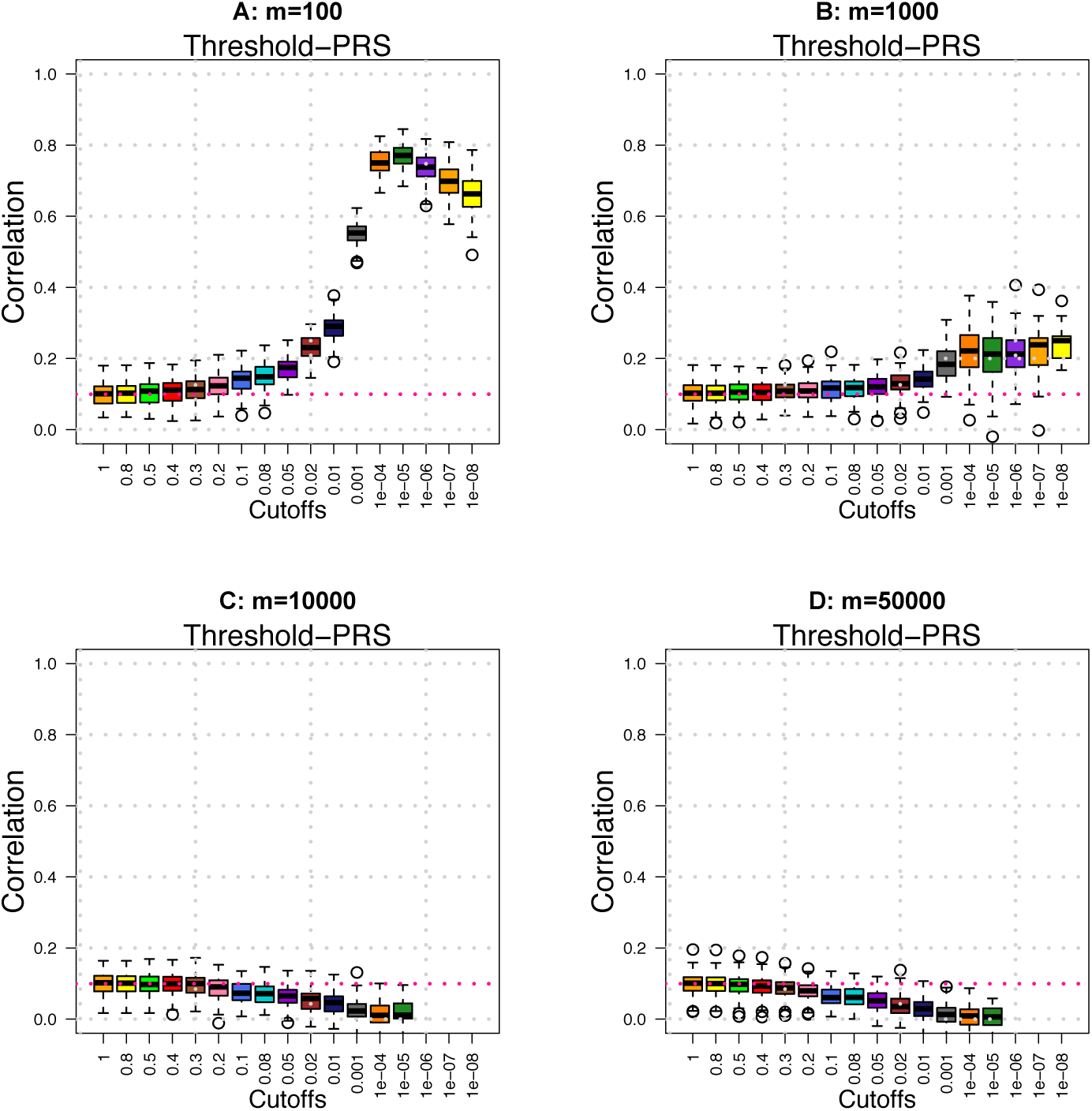
Prediction accuracy (*A*_*P*_) of threshold-PRS across different *m/p* ratios when the effects of causal SNPs are not i.i.d. Normal. SNP data are independently sampled from {0, 1, 2}. We set *p*=100, 000 and *n*=1000 in both training and testing data.

**Supplementary Figure 11:**
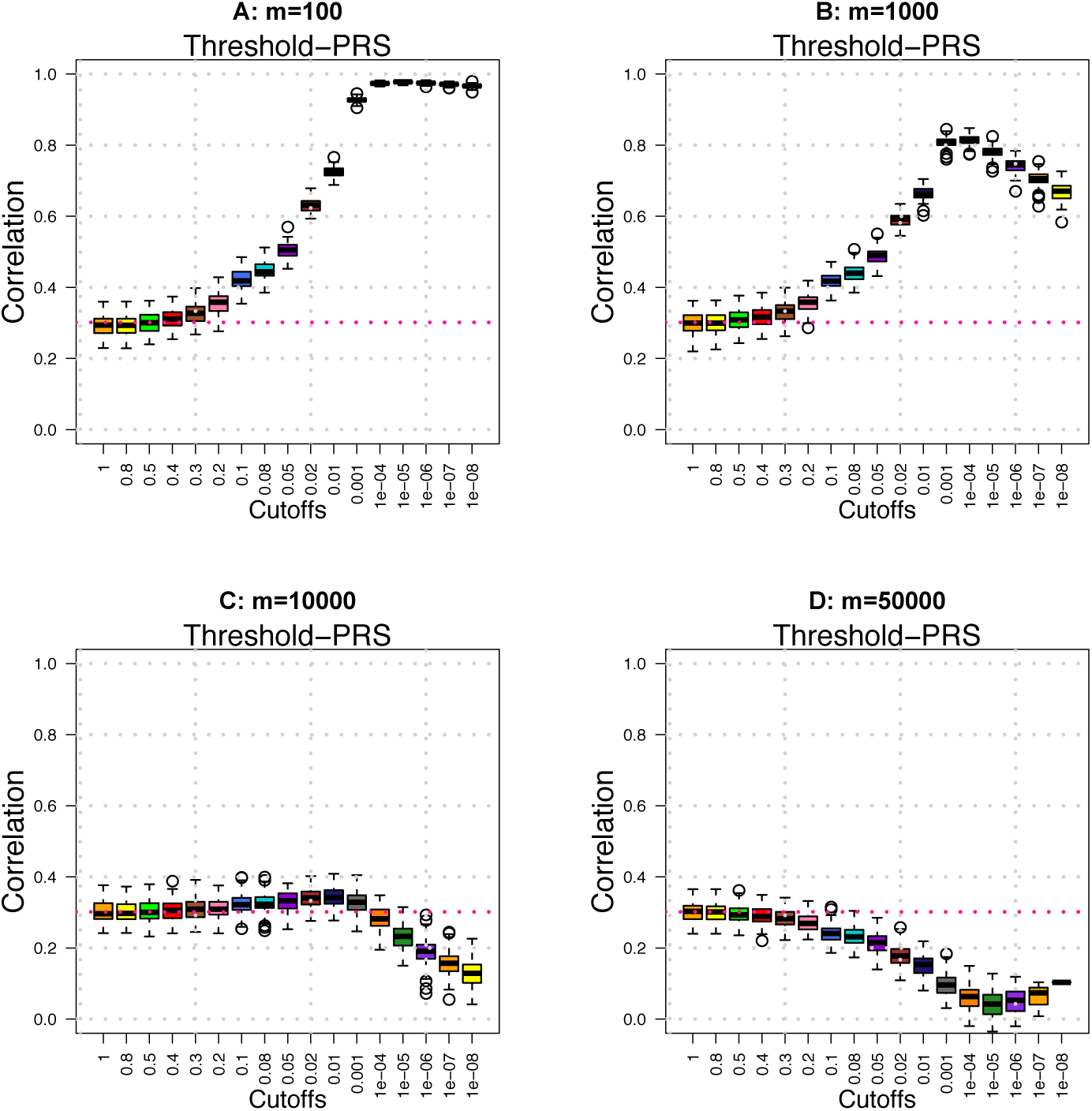
Prediction accuracy (*A*_*P*_) of threshold-PRS across different *m/p* ratios when the effects of causal SNPs are not i.i.d. Normal. SNP data are independently sampled from {0, 1, 2}. We set *p*=100, 000 and *n*=10, 000 in training data and *n* = 1000 in testing data.

**Supplementary Figure 12:**
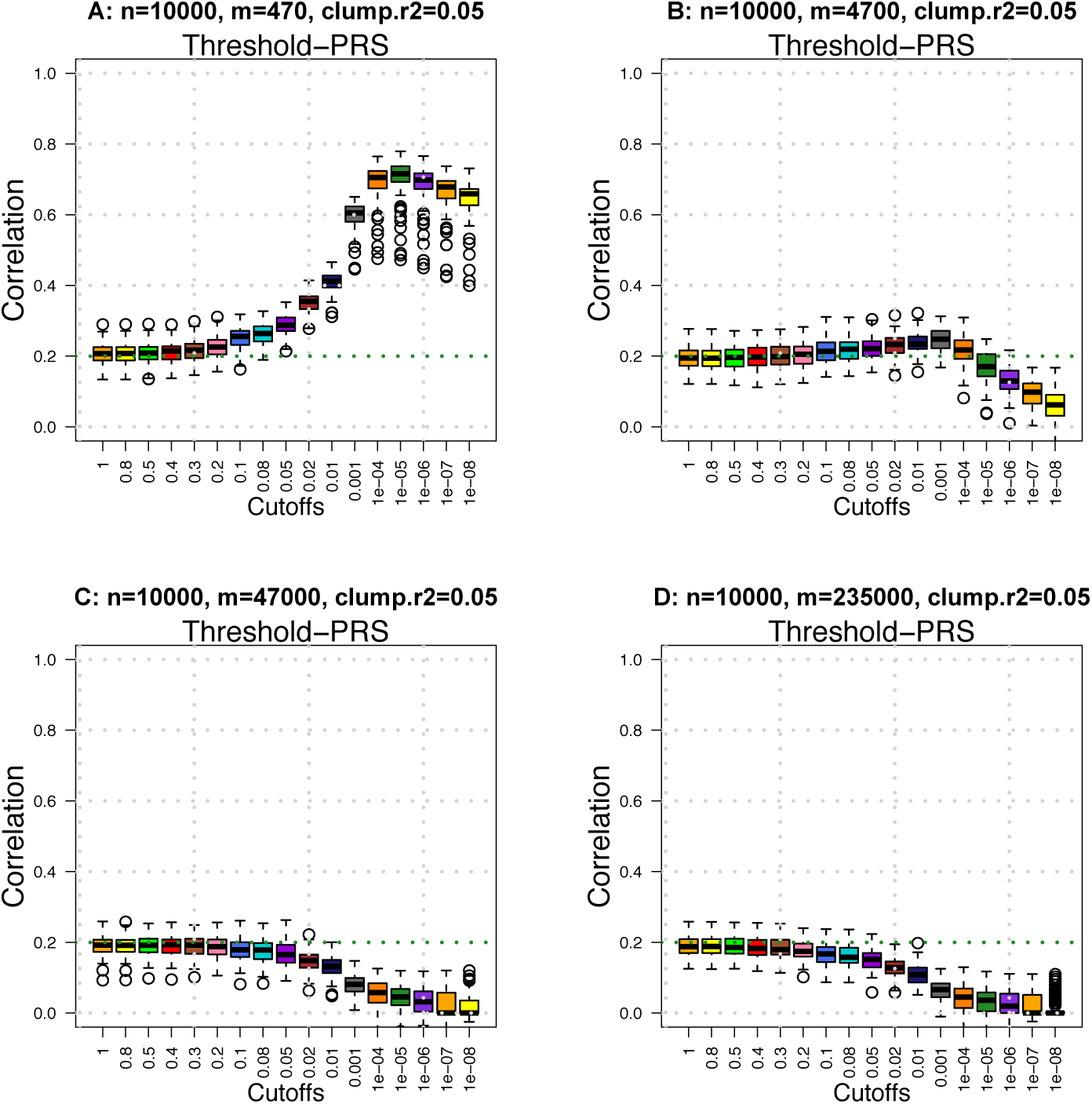
Prediction accuracy (*A*_*P*_) of threshold-PRS across different *m/p* ratios in the UKB data simulation. There are *p* = 461, 488 SNPs and *n* = 10, 000 individuals in the training data. Clumping *R*^2^ parameter is set to be 0.05. *p* ≈ 150, 000 SNPs pass LD-based clumping and are candidates SNPs to construct PRS on *n* = 1000 subjects in testing data.

**Supplementary Figure 13:**
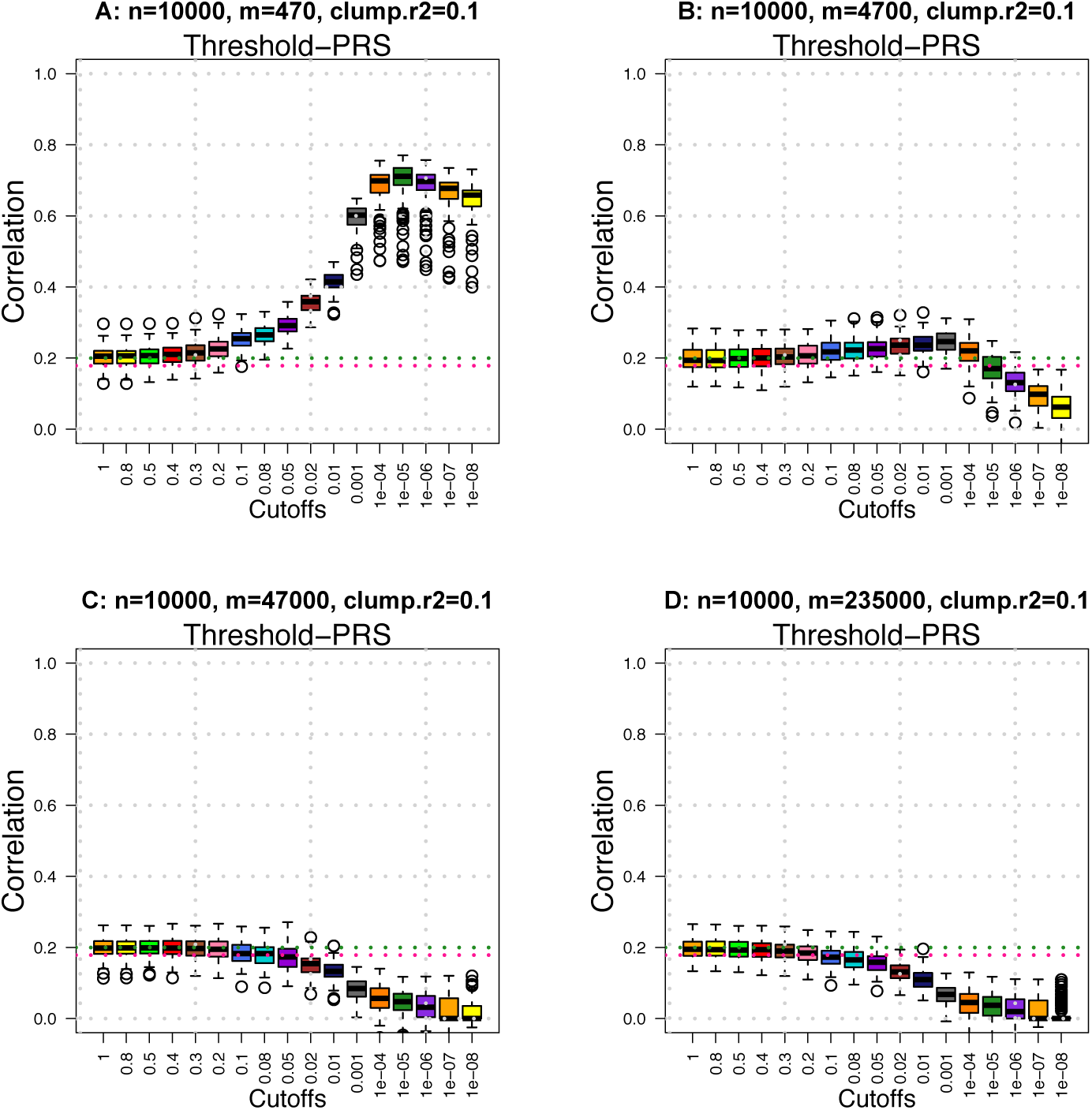
Prediction accuracy (*A*_*P*_) of threshold-PRS across different *m/p* ratios in the UKB data simulation. There are *p* = 461, 488 SNPs and *n* = 10, 000 individuals in the training data. Clumping *R*^2^ parameter is set to be 0.1. *p* ≈ 190, 000 SNPs pass LD-based clumping and are candidates SNPs to construct PRS on *n* = 1000 subjects in testing data.

**Supplementary Figure 14:**
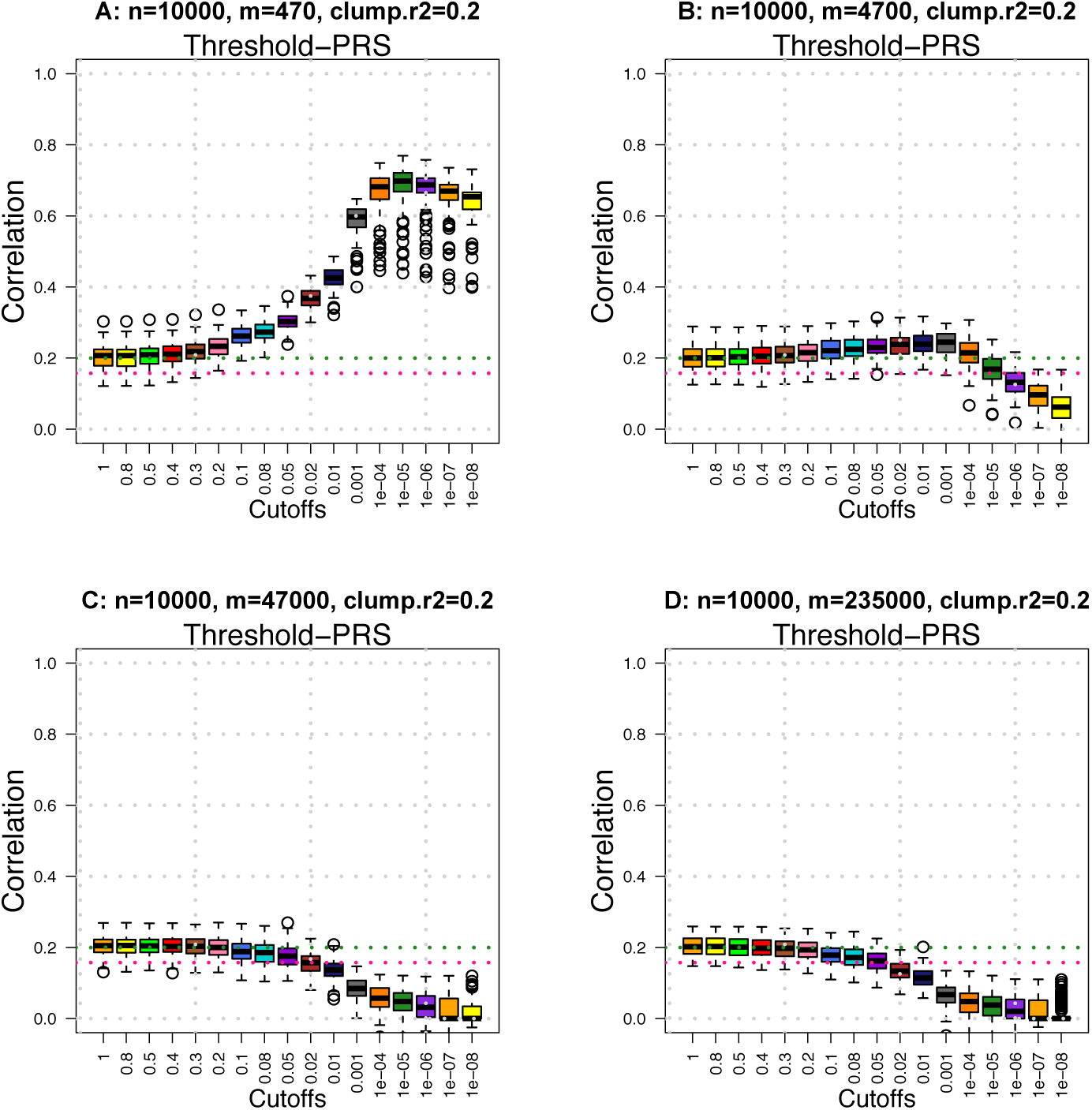
Prediction accuracy (*A*_*P*_) of threshold-PRS across different *m/p* ratios in the UKB data simulation. There are *p* = 461, 488 SNPs and *n* = 10, 000 individuals in the training data. Clumping *R*^2^ parameter is set to be 0.2. *p* ≈ 250, 000 SNPs pass LD-based clumping and are candidates SNPs to construct PRS on *n* = 1000 subjects in testing data.

**Supplementary Figure 15:**
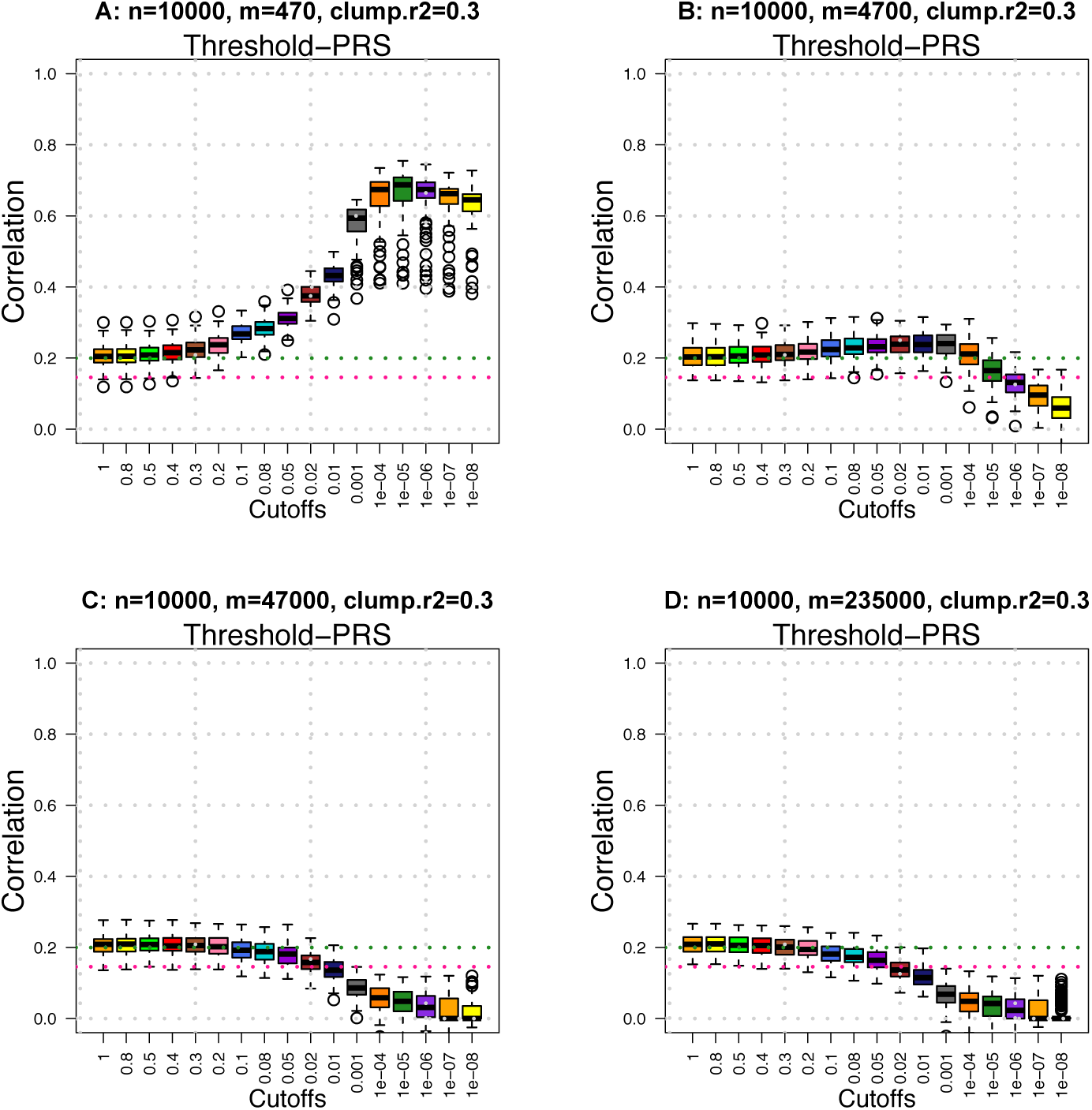
Prediction accuracy (*A*_*P*_) of threshold-PRS across different *m/p* ratios in the UKB data simulation. There are *p* = 461, 488 SNPs and *n* = 10, 000 individuals in the training data. Clumping *R*^2^ parameter is set to be 0.3. *p* ≈ 290, 000 SNPs pass LD-based clumping and are candidates SNPs to construct PRS on *n* = 1000 subjects in testing data.

**Supplementary Figure 16:**
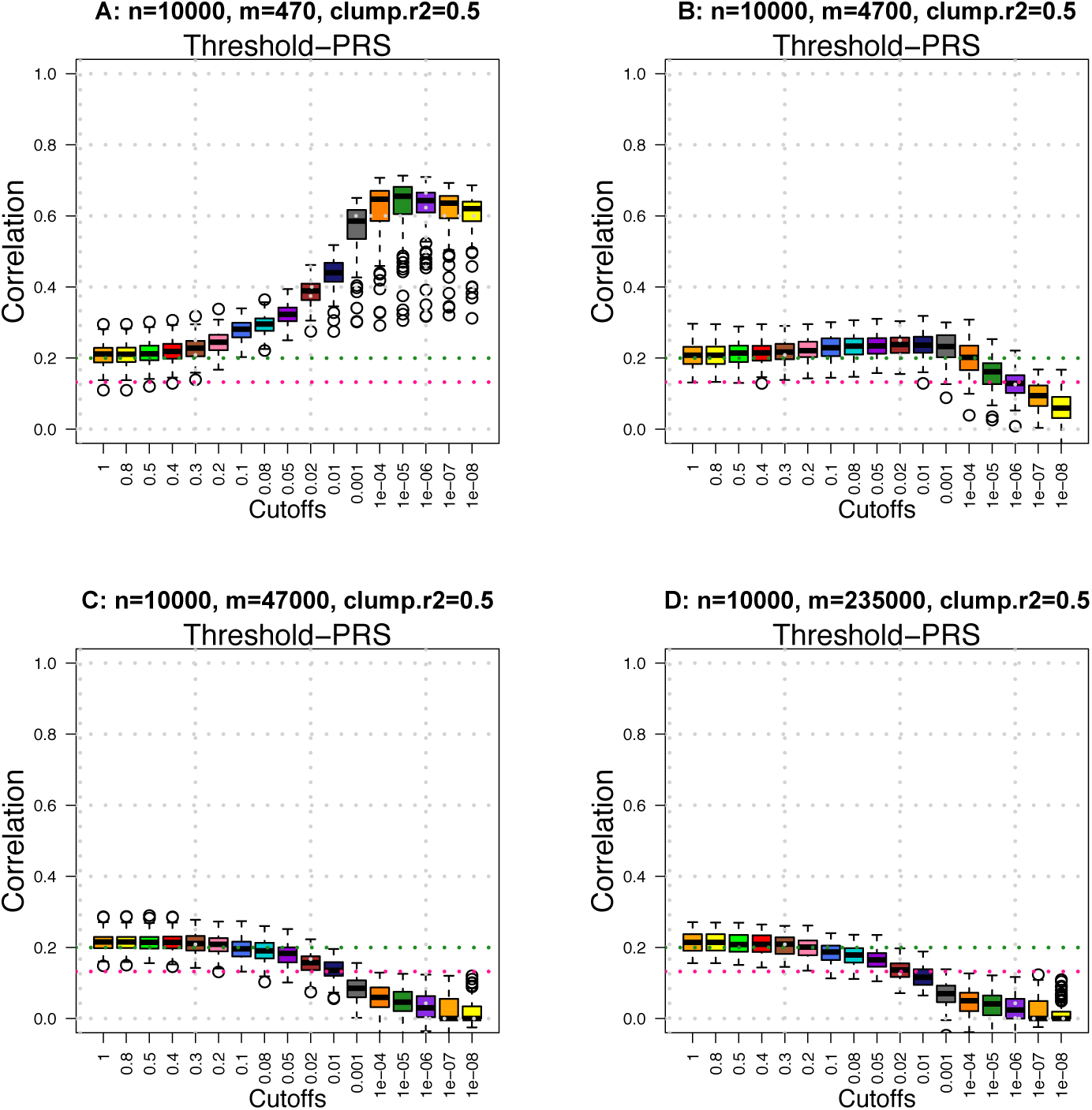
Prediction accuracy (*A*_*P*_) of threshold-PRS across different *m/p* ratios in the UKB data simulation. There are *p* = 461, 488 SNPs and *n* = 10, 000 individuals in the training data. Clumping *R*^2^ parameter is set to be 0.5. *p* ≈ 360, 000 SNPs pass LD-based clumping and are candidates SNPs to construct PRS on *n* = 1000 subjects in testing data.

**Supplementary Figure 17:**
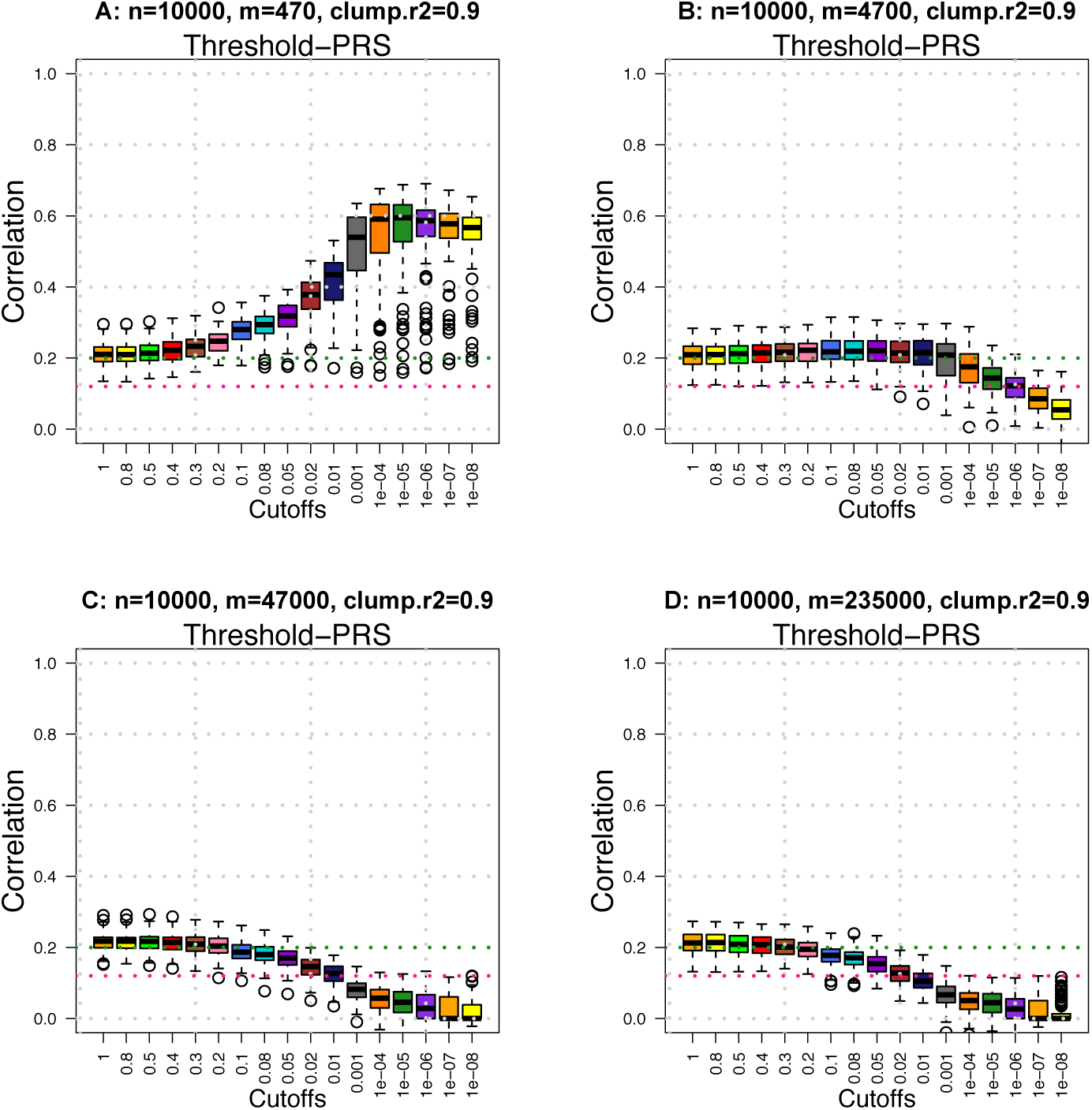
Prediction accuracy (*A*_*P*_) of threshold-PRS across different *m/p* ratios in the UKB data simulation. There are *p* = 461, 488 SNPs and *n* = 10, 000 individuals in the training data. Clumping *R*^2^ parameter is set to be 0.9. *p ≈* 430, 000 SNPs pass LD-based clumping and are candidates SNPs to construct PRS on *n* = 1000 subjects in testing data.

**Supplementary Figure 18:**
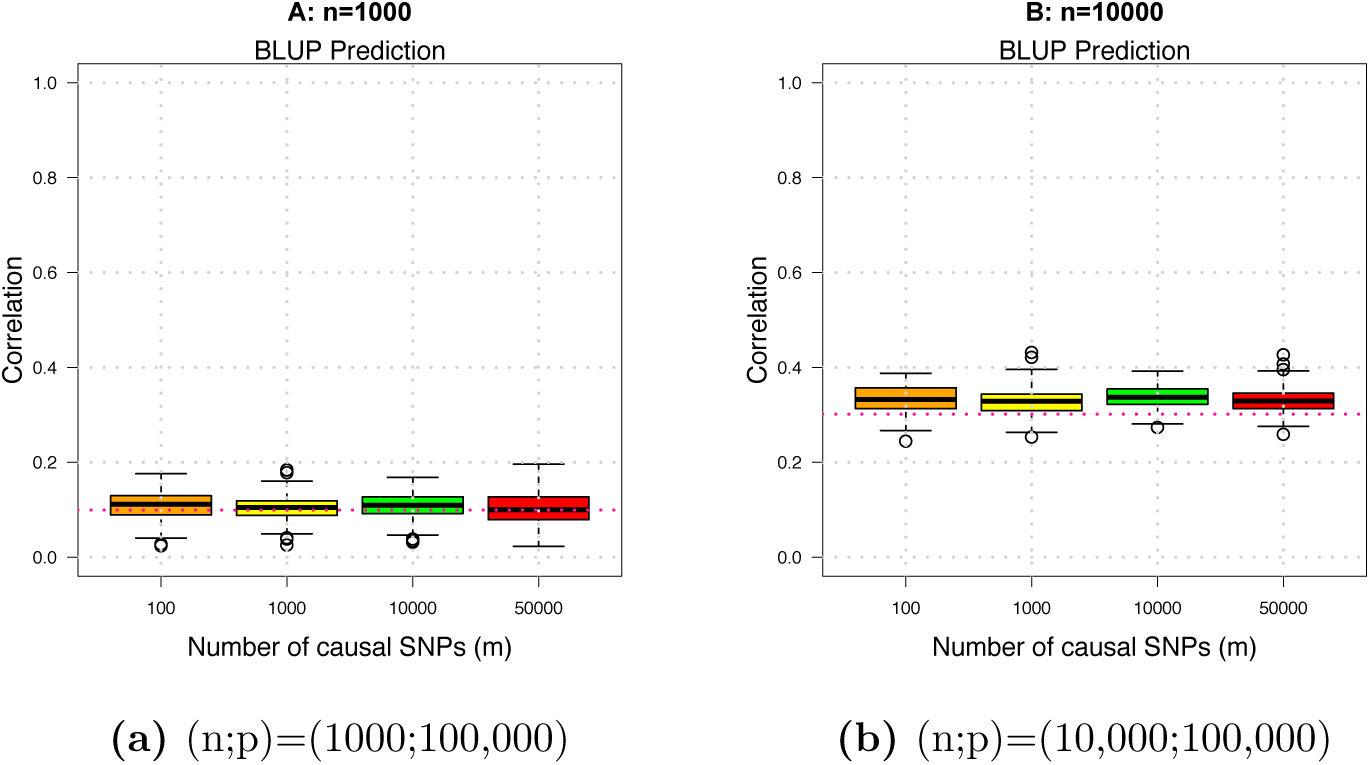
Prediction accuracy (*A*_*P*_) of BLUP across different *m/p* ratios. We set *p*=100, 000 and *m* to be 100, 1000, 10, 000, and 50, 000, respectively. In (a), we set *n*=1000 in both training and testing data, and in (b), we set *n*=10, 000 in training data and *n*=1000 in testing data.

**Supplementary Figure 19:**
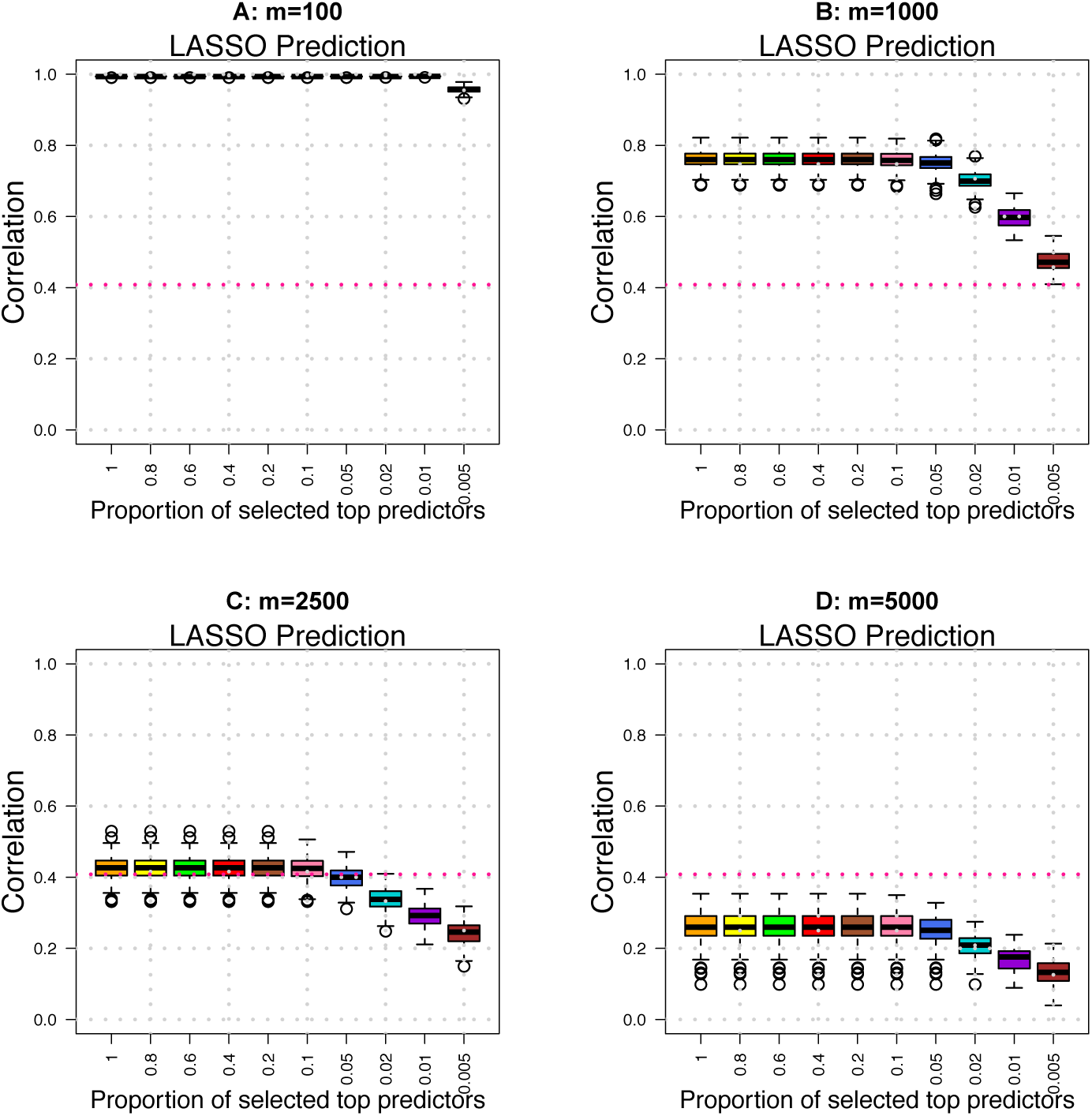
Prediction accuracy (*A*_*P*_) of LASSO across different *m/p* ratios. We set *p*=10, 000 and *n*=2000 in training data and *n*=1000 in testing data.

**Supplementary Figure 20:**
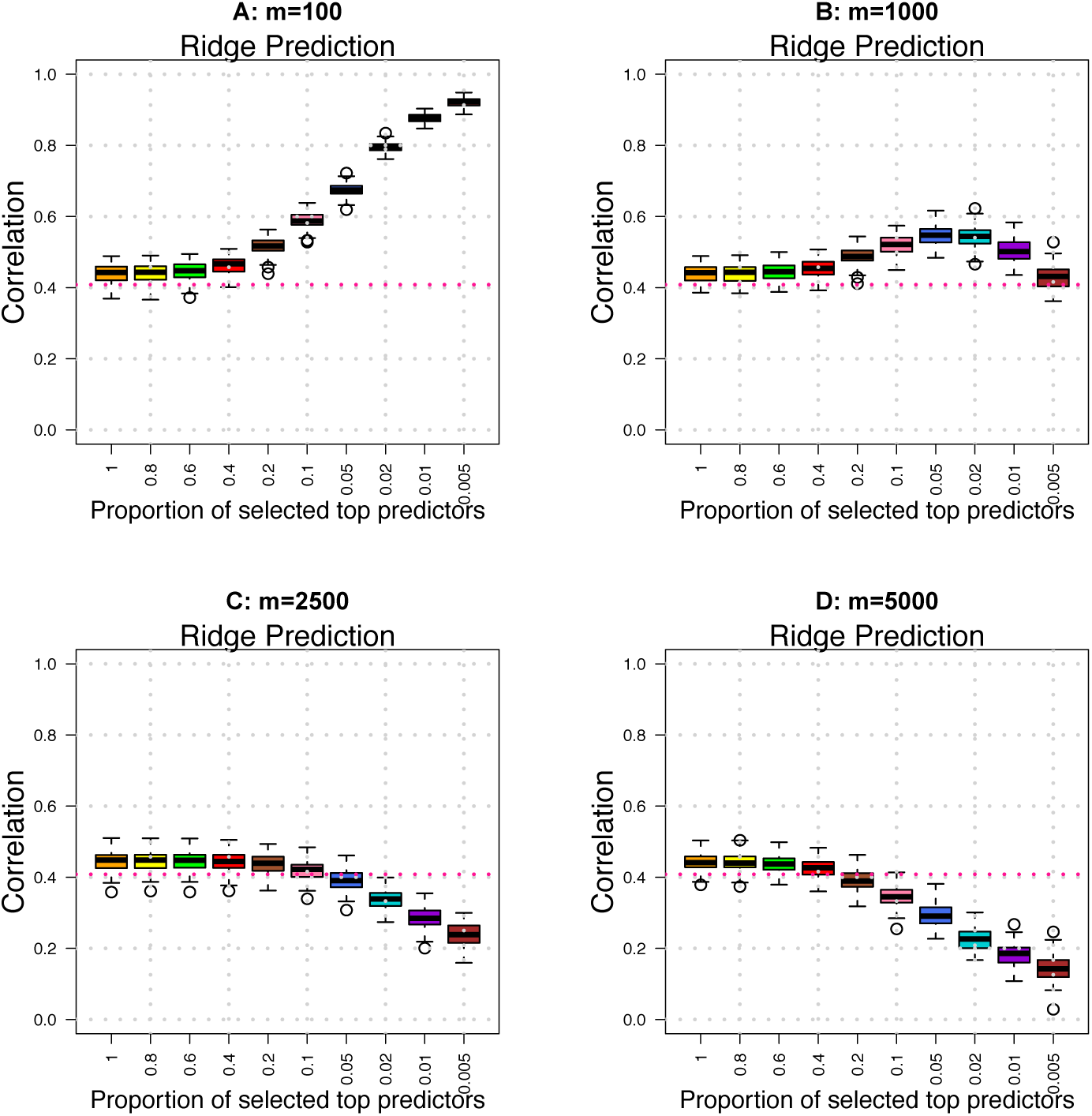
Prediction accuracy (*A*_*P*_) of ridge regression across different *m/p* ratios. We set *p*=10, 000 and *n*=2000 in training data and *n*=1000 in testing data.

